# An assessment of phylogenetic tools for analyzing the interplay between interspecific interactions and phenotypic evolution

**DOI:** 10.1101/083485

**Authors:** J.P. Drury, Grether G.F., T. Garland, H. Morlon

## Abstract

Much ecological and evolutionary theory predicts that interspecific interactions often drive phenotypic diversification and that species phenotypes in turn influence species interactions. Several phylogenetic comparative methods have been developed to assess the importance of such processes in nature; however, the statistical properties of these methods have gone largely untested. Focusing mainly on scenarios of competition between closely-related species, we assess the performance of available comparative approaches for analyzing the interplay between interspecific interactions and species phenotypes. We find that currently used statistical methods largely fail to detect the impact of interspecific interactions on trait evolution, that sister taxa analyses often erroneously detect character displacement where it does not exist, and that recently developed process-based models have more satisfactory statistical properties. In weighing the strengths and weaknesses of different approaches, we hope to provide a clear guide for empiricists testing hypotheses about the reciprocal effect of interspecific interactions and species phenotypes and to inspire further development of process-based models.

---

Interactions between species are a fundamental aspect of life on earth, and understanding the evolutionary and ecological consequences of such interactions are a central goal of many classical theoretical frameworks in ecology and evolutionary biology. Identifying both the predictors of interspecific interactions and the consequences of such interactions for diversification and coexistence is thus an important contemporary research area, with strong implications for conservation biology.

Several phylogenetic comparative methods have been deployed with the goal of elucidating how interspecific interactions drive (or are driven by) character evolution, but the reliability and efficacy of these methods remain largely untested. Here we focus on methods used to study interactions between closely related species (e.g., members of the same family) that arise from similarity in morphology, signaling traits or habitat (Brown and Wilson 1956; Schluter 2000; Pfennig and Pfennig 2009), rather than on community-wide interactions and interaction networks (Webb et al. 2002; Rezende et al. 2007; Cavender-Bares et al. 2009; Cadotte et al. 2013).

Classical character displacement theory (Brown and Wilson 1956; Grether et al. 2009; Pfennig and Pfennig 2009) predicts that, where heterospecifics compete, selection should favor divergence in the traits responsible for competition, until lineages in sympatry no longer compete intensely. In a seminal example, selection resulting from exploitative competition between medium and large ground finches (*Geospiza fortis* & *G. magnirostris)* has driven bill size divergence on Daphne Major in the Galápagos (Grant and Grant 2006). Investigators who conduct comparative studies of divergent character displacement often test for a relationship between biogeographic overlap and trait dissimilarity, predicting that coexisting species will be more phenotypically divergent than non-coexisting ones. Recent studies on *Bicyclus* butterflies and *Euglossa* bees, for example, show that male chemical cues are more distinct between sympatric species than allopatric species, suggesting that reproductive character displacement has driven signal divergence in these taxa (Bacquet et al. 2015; Weber et al. 2016).

Interspecific interactions can also lead to convergent, rather than divergent, character displacement (Cody 1969, 1973;Grant 1972; Grether et al. 2013). Agonistic character displacement theory (Grether et al. 2013) predicts convergence in traits mediating interspecific aggression when species compete strongly for the same resources. In other words, between-species similarity in resource use may make interspecific territoriality adaptive, resulting in subsequent convergence in signaling traits involved in mediating territorial interactions (e.g., song in ovenbirds, Tobias *et al.* 2014). Therefore, tests of convergent character displacement typically test the prediction that sympatric lineages are more phenotypically similar than allopatric ones. Because sympatric similarity can also result from convergence to local conditions (e.g., habitat, climate), it is important for empiricists to account for abiotic factors in tests of character convergence.

In some instances, rather than identifying the effect of species interactions on trait evolution, empiricists aim to identify traits that mediate particular pairwise interactions, such as hybridization or interspecific aggression. In this case, investigators test for a relationship between the measured interactions and trait similarity. Recent studies on New World warblers (Parulidae), for example, show that hybridization occurs more often between species with similar songs and that interspecific territoriality occurs more often between species that share similar plumage and territorial song phenotypes (Willis et al. 2014; Losin et al. 2016).

Although the examples presented here largely represent scenarios where interactions between species are competitive, empiricists apply methods discussed here to other non-competitive interactions as well (e.g., predicting links in plant/pollinator networks, identifying Müllerian mimicry rings: Elias et al. 2008; Eklöf and Stouffer 2016). Regardless of the biological question, a particularity of comparative tests aimed at understanding the interplay between interspecific interactions and species phenotypes is that they largely involve testing correlations between pairwise data (e.g. range overlap, phenotypic similarity, frequency of hybridization). In contrast, most phylogenetic comparative methods have been developed and tested on tip data (e.g. range size, morphological trait values), and the statistical properties of methods adapted to handle pairwise data (Box 1) have gone untested (but see Harmon & Glor 2010). Furthermore, species interactions are inherently affected by the biogeographic history of dispersal and speciation in an evolving clade and the resulting patterns of range overlap. Patterns of trait dissimilarity between sympatric lineages—the classic test of character displacement— may actually be the null expectation if allopatric speciation is the norm, because then sympatric species pairs will tend to share more distant common ancestors than allopatric species pairs do (Weir and Price 2011; Tobias et al. 2014).

Here, we apply the main phylogenetic comparative methods that investigators use to test hypotheses about interactions between closely related lineages and phenotypes (Box 1, Fig. 1) to datasets simulated under different evolutionary histories of speciation, dispersal, species interactions, and trait evolution. We then compare the efficacy of these methods, discuss the relative merits of each, and outline directions for future research.

**Figure 1.**
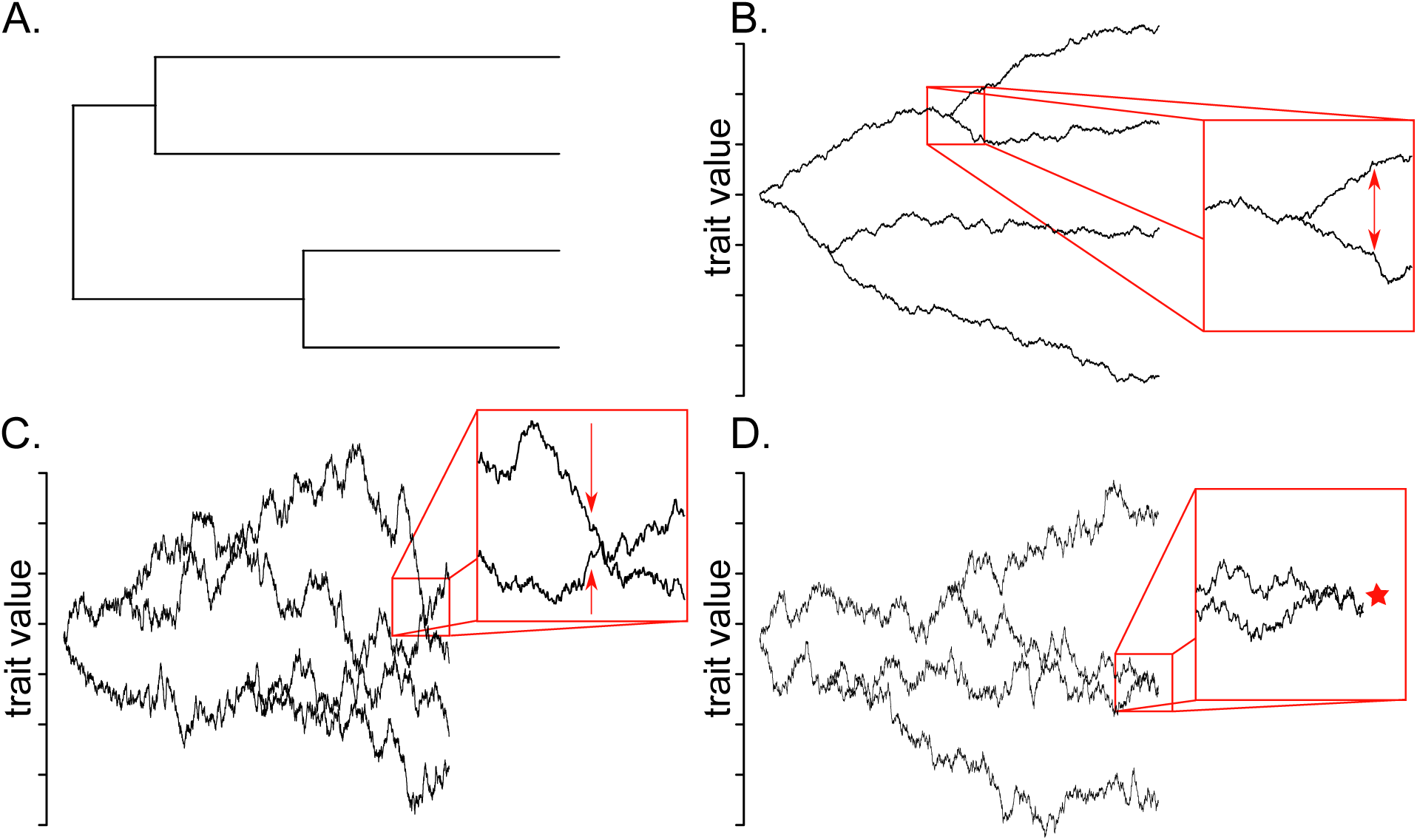
Schematic examples of the processes examined in our simulation study. A. Phylogeny along which the trait evolves. B. A trait evolving via divergent character displacement, C. A trait evolving via convergent character displacement, and D. A species interaction that exists at present due to pairwise trait similarity. For simulation details, see the main text and *Supplementary Methods*.

## METHODS

We compared the performance of different phylogenetic comparative methods by measuring their statistical power (e.g., probability of detecting divergence when divergence is simulated) and Type I error rate (e.g., probability of detecting an effect of species interactions when such an effect is not simulated) across three scenarios.

### Phylogeny and Range Simulations

We jointly simulated trees {# spp. = 20, 50, 100, 150, 200, 250} and biogeographies under the dispersal-extinction-cladogenesis model of biogeographical evolution (i.e., DEC+J, with the inclusion of founder event speciation) in BioGeoBEARS (Ree and Smith 2008; Matzke 2014). Briefly, the DEC+J model is a model of range evolution in which species ranges change along the branches of a phylogeny as a function of dispersal and local extinction and are inherited by daughter taxa at speciation according to several possible cladogenetic scenarios (see more details in Supplementary Methods). For each tree, we started with a single ancestral species occupying one of ten equidistant regions, and simulated trees with constant rates of speciation and local extinction. We considered different biogeographic scenarios by varying the rate of dispersal events between ranges (“high” and “low” dispersal; see details in Supplementary Methods) and the probability that speciation events occur in sympatry versus allopatry (“high” and “low” sympatric speciation; Supplementary Methods). Each of these simulations resulted in a phylogeny (the tree of extant species) and its associated biogeography (the set of regions in which each lineage occurred throughout the history of the clade). Lineages were identified as sympatric if they co-occurred in at least one of the ten geographic regions, and allopatric if they did not co-occur in any.

We simulated four biogeographic scenarios (combinations of low or high dispersal and low or high sympatric speciation) for each tree size. The resulting biogeographies span scenarios where sympatric speciation is common and dispersal is low (e.g., lizards on islands) to scenarios where allopatric speciation is the main mode of speciation and dispersal between regions is high (e.g., birds on continents). These parameter combinations produced a range of realistic proportions of sister taxa that are sympatric (Fig. S1A) and a range of realistic differences in age between sympatric and allopatric sister taxa (Pigot and Tobias 2014; Fig. S1B). In defining sympatry as any overlap, the mean magnitude of range overlap fell between 33-42% across all tree sizes and simulation parameters (Fig. S1C,D), which falls well within the range of overlap of sympatric taxa defined under commonly used minimum threshold values applied to continuous indices of range overlap (e.g. Pigot and Tobias 2014; Tobias et al. 2014).

For each combination of tree sizes and DEC parameter combinations (*n* = 24), we performed 100 simulations, resulting in a bank of 2,400 trees with associated biogeographies.

### Character Displacement

#### The model

To simulate both divergent and convergent character displacement, we simulated a continuous trait *z* under a model in which trait values of sympatric species in an evolving clade are repelled from (or drawn toward) one another. In divergent character displacement, trait divergence is driven by pairwise similarity in that same trait *z*; in convergent character displacement however, convergence in trait z (e.g. a signaling trait) is driven by pairwise similarity in another trait *y* (e.g. a resource use trait). To create a generic model of character displacement, we thus modified the matching competition model (Nuismer & Harmon 2015; Drury *et al.* 2016) by describing the mean value for trait *z* in lineage *i* after an infinitesimally small time step *dt* by:

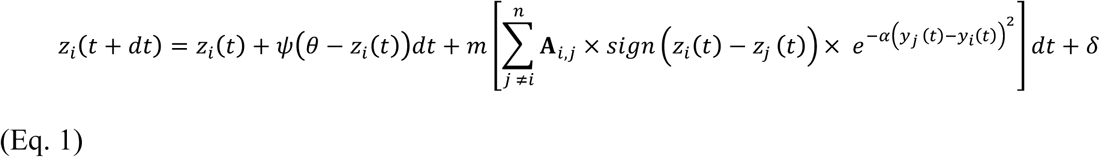

where *y* = z in the case of divergent character displacement and *y* ≠ *z* in the case of divergent character displacement, *Ψ* (θ − *z_i_*(*t*)) describes attraction to a single stationary peak (i.e., the Ornstein-Uhlenbeck [OU] process, Felsenstein 1988; Garland et al. 1993; Hansen and Martins 1996), *n* is the number of species, *δ* is a random variable with mean 0 and variance = *σ*^2^ *dt* (the Brownian motion [BM] rate parameter, describing the stochastic component of trait evolution), and **A** is a piecewise-constant matrix representing biogeographical overlap such that **A**_*i,j*_ equals 1 if species *i* and *j* are sympatric at time t, and 0 otherwise. The “sign” portion determines the relative position of each species in trait space (i.e. it equals +1 if *z_i_* is larger than *z_j_,* and −1 otherwise). The *α* value (*α* > 0) determines the effect of pairwise similarity in trait *y* on competition: if α is close to zero, all lineages sympatric with lineage *i* have the same competitive effect on *i*, regardless of their similarity in trait *y*; conversely, if *α* is large, sympatric lineages similar to i in terms of the *y* trait will have a much stronger competitive effect on *i* than sympatric lineages dissimilar to i in terms of the *y* trait. The parameter *m* represents the magnitude of the effect of competition when two lineages have identical *y* values (i.e., it provides an upper bound for the deterministic effect of competition). When *m* = 0, this equation reduces to an OU model, whereas positive *m* values result in pairwise divergence and negative values result in pairwise convergence. When both *m* and *ψ =* 0, this model reduces to Brownian motion. For additional simulation details, see *Supplementary Methods.*

We use a lineage-based “phenomenological” model for our simulations rather than an individual-based model to have the computational ability to produce datasets of a size comparable to the maximum sometimes reached in empirical comparative phylogenetic studies (i.e. often reaching several hundreds of species). Models derived from microevolutionary first principles (e.g., Grether et al. 2009; Nuismer and Harmon 2015) generate similar patterns of sympatric shifts resulting from character displacement, and using such a model here would be much more computationally intensive, therefore restricting the range of parameter values that can be studied. For simplicity, this model also omits the effect of a species’ geographic structure and the effect of gene flow between distinct populations on the evolution of the mean species phenotype. This simplification is reasonable in the context of our study because there is no reason to expect that it will systematically bias the patterns generated in such a way as to yield different conclusions regarding the performance of the various analytical approaches that we use here. Finally, in all of our simulations, we considered sympatry to be a binomial variable, so **A**_i,j_ equaled either 1 (if species *i* and *j* are sympatric) or 0 (if species *i* and *j* are allopatric). This index of sympatry is similar to commonly used indices (Pigot and Tobias 2014; Tobias et al. 2014), but other formulations of sympatry, such as continuous measurements of range overlap (Bothwell et al. 2015; Martin et al. 2015) are also possible. We did not explore continuous measurements of range overlap here, but have uploaded our simulation scripts to RPANDA (Morlon et al. 2016; https://github.com/hmorlon/PANDA), which could easily be modified to do so.

#### Divergent character displacement

We simulated datasets with divergent character displacement by setting *y* = *z* in Eq. 1 such that trait divergence is driven by pairwise similarity in that trait. Biologically, this could represent a feeding trait that co-varies with resource use (e.g., bill shape in Galápagos finches, Grant & Grant 2011) and which would be directly affected by interspecific competition. To assess whether each method could detect divergent character displacement when it occurred and did not erroneously detect character displacement when it was absent, we simulated datasets both with repulsion {*m* = 2} and without repulsion {*m* = 0} (see Supplementary Methods). We also simulated datasets with {*ψ* = 2} and without {*ψ* = 0} the OU process. In all simulations, we held *σ*^2^ constant at 0.5, *α* constant at 1, and both the state at the root (*z*_0_) and the OU optimum (*θ*) constant at 0.

In additional simulations run only on 100-species trees, we analyzed the effect of both the maximum strength of repulsion {*m* = 0, 1, 2, 10} and, to understand how the opposing forces of repulsion and attraction to an optimum influence analyses, the ratio of attraction to the maximum effect of competition {*ψ:m* = 0, 0.2, 0.5, 1} on inferences. To achieve these ratios of *ψ:m,* we varied *ψ* while holding *m* constant (e.g., for the case where *m* = 2, we simulated datasets where *ψ* = 0, 0.4, 1, and 2, respectively). As above, these values were arbitrarily chosen based on visual inspection of realized simulations.

For each parameter combination, we simulated 10 datasets for each tree, resulting in 1,000 simulations for each tree size / biogeographic scenario combination.

#### Convergent character displacement

We simulated datasets with convergent character displacement under Eq. 1, where the term *y* represents a trait determining resource use or niche occupation evolving via BM or OU. A species’ trait *z* in this model—a trait used as a territorial signal—is thus attracted most strongly to the signal trait values of sympatric lineages with the most similar resource-use traits. Biologically, this represents a scenario where selection favors interspecific territoriality—mediated by similarity in territorial signals—because the benefits of excluding heterospecifics are similar to the benefits of excluding conspecifics (Grether et al. 2009). As a species’ resource-use trait becomes less similar to that of sympatric species, the strength of attraction decreases to zero.

We simulated resource-use traits under both BM (*σ*^2^_*resource*_ = 0.5, *ψ*_*resource*_ = 0) and OU (*σ*^2^_*resource*_ = 0.5, *ψ_resource_* = 2, *θ_resource_* = 0) models. For the signal trait, we simulated datasets both with convergence {*m* = −0.25} and without convergence {*m* = 0}. We did not include attraction toward a stable peak for the signal trait (i.e. *ψ* was held constant at 0). As above, we held *σ*^2^ = 0.5 and *z*_0_ = 0, though we held *α* constant at 10, since smaller values result in rapid, cladewise convergence in traits. To analyze the effect of the maximum strength of convergence, we ran another set of simulations on 100-species trees varying *m* {*m* = 0, −0.1, −0.25, −0.5} (see Supplementary Methods). The resource trait (*y*) and signal trait (*z*) were modeled as unlinked and genetically uncorrelated.

As above, we simulated 10 datasets for each tree, resulting in 1,000 simulations for each tree size / biogeographic scenario combination.

### Predictors of Interspecific Interactions

In some cases, investigators wish to identify which factors explain the occurrence of particular interspecific interactions. For example, investigators may want to understand which traits cause species to hybridize (e.g., Willis et al. 2014). In this scenario, species interactions vary according to phenotypic similarity between sympatric species pairs (i.e., species pairs that could potentially interact). Additionally, and unlike character displacement analyses, predicting the occurrence of interspecific interactions requires treating trait similarity as a predictor variable rather than a response variable. Thus, we generated datasets where the presence of interactions between sympatric taxa depends on pairwise similarity in traits.

Under this scenario, we first evolved a trait along the phylogeny under a BM (*σ*^2^ = 0.5, *ψ* = 0) or OU (*σ*^2^ = 0.5, *ψ* = 2, *θ* = 0) model. Next, we simulated a second, independently evolving trait (*σ*^2^_*unmeasured*_ = 1, *ψ_unmeasured_* = 0) to represent an unmeasured trait that could cause the interaction of interest. To generate datasets where species interactions depend on similarity in trait space at the present, we created species interactions in the form of a binomial variable by sampling from a binomial distribution with the probability of interaction equal to:

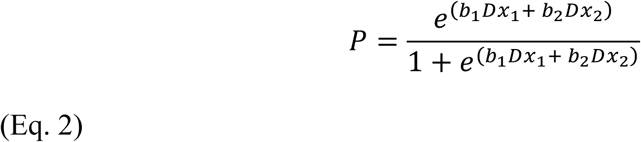

(e.g., Hilbe 2009) where *Dx_n_* is the distance between species at the present (e.g., distance between tip values) in simulated trait *n* (simulated using fastBM in phytools, Revell 2012), and *b_n_* is the coefficient determining the magnitude of the relationship between the species interaction and similarity in trait *n*. Trait 1 is the measured, focal trait and trait 2 represents the independently evolving, unmeasured trait. As the effect of *b_n_* on the species interaction depends on the *Dx_n_* distribution, which in turn depends on the total height of the tree, we scaled the trees to a height of one prior to simulating datasets to facilitate comparison of results across trees and parameter space.

To determine the statistical power of each analytical method, we generated species interactions based on similarity in the measured trait (*b_1_* = −4, *b_2_* = 0); to assess the Type I error rate, we simulated species interactions based on similarity in the unmeasured trait (*b_1_* = 0, *b_2_* = −4). To determine the effect of the magnitude of the coefficient determining the relationship between the measured trait and the interactions, we ran another set of simulations on 100-species trees varying *b_1_* {*b_1_* = 0, −2, −4, −6, −8} and holding *b_2_* = −4. As above, we ran 1,000 simulations for each tree size / biogeographic scenario combination.

### Phylogenetic Tests

Among our tests of character displacement (both divergent and convergent), the “correlation” tests involved assessing the significance of the relationship between phenotypic similarity and coexistence, using either the “full” dataset (all species pairs) or the “sister taxa” subset obtained by culling sister taxa from trees with ≥150 tips (Box 1, Diagram S1). To the full datasets, we applied standard non-phylogenetic regression analyses that ignore phylogenetic nonindependence (Box 1.1), the raw and phylogenetically permuted partial Mantel tests (Box 1.2, 1.3), phylogenetic linear mixed models (PLMMs, Box 1.4), and the simulation approach (Box 1.5, Supplementary Methods). To the sister-taxa datasets, we applied non-phylogenetic regression analyses (Box 1.1), PLMMs (Box 1.4), the simulation approach (Box 1.5), sister-taxa GLMs (Box 1.7), and fit process based models in EvoRAG (Box 1.8, Supplementary Methods). We did not perform Mantel tests on the sister-taxa data because such tests require complete matrices and distance matrices with data for only sister taxa would mostly contain empty cells (i.e. all those cells that correspond to non sister taxa species pairs). We compared the fit of process-based phenotypic models with and without species interactions (Brownian motion, Ornstein-Uhlenbeck, diversity dependent, and matching competition models; see Box 1.6 and Supplementary Methods) to the full datasets from divergence scenarios using the R packages geiger (Pennell et al. 2014) and RPANDA (Morlon et al. 2016). We acknowledge that diversity-dependent models were not designed to analyze character displacement *per se,* but because they incorporate interspecific interactions, we hypothesized that (and wanted to test if) they could be useful in doing so. We did not apply process-based models to convergence scenarios because the necessary model fitting tools have yet to be developed (see Discussion).

Our tests of predictors of species interactions involved assessing the significance of the relationship between phenotypic similarity and species interactions (i.e., whether the species interact where they occur in sympatry). Since the response variable is binary, we fit non-phylogenetic logistic regressions, logistic PLMMs, and employed the simulation approach (see Supplementary Methods). We did not perform Mantel tests or sister-taxa analyses because the species pair matrix was incomplete (species that do not coexist cannot interact) and typically too few sister taxa occurred in sympatry for regression analysis.

## RESULTS

### Divergent Character Displacement

When all possible pairwise comparisons are included in analyses, the ability of most methods to detect divergent character displacement depends on the presence of the OU process. As expected, non-phylogenetic regression analyses have a high Type I error rate in either scenario (Figs. 2Ai,iv, S2Ai,iv [NB: throughout, results for low sympatric speciation biogeographies are plotted in the main text and high sympatric speciation biogeographies in the supplement]). When the OU process is present (*ψ* = 2), all methods generally have low Type I error rates and high power (Figs. 2Aiv-vi, S2iv-vi, Supplementary Tables). However, when there is no pull toward a peak (*ψ* = 0), the Type I error rate is higher for Mantel tests (Figs. 2Ai-ii, S2i-ii), and the power is much lower for all methods, though the pppMantel and raw Mantel perform better than the simulation and PLMM methods (Figs. 2Aiii, Fig. S2iii). Repulsion is easier to detect against an OU background of traits converging toward a common optimum than against a background of traits diverging under BM, likely because the repulsion process is more active when species occupy similar trait space (Figs. S3, S4). High rates of sympatric speciation and dispersal tend to slightly decrease the power of all methods (Fig. S2ii,iv, Supplementary Tables).

**Figure 2.**
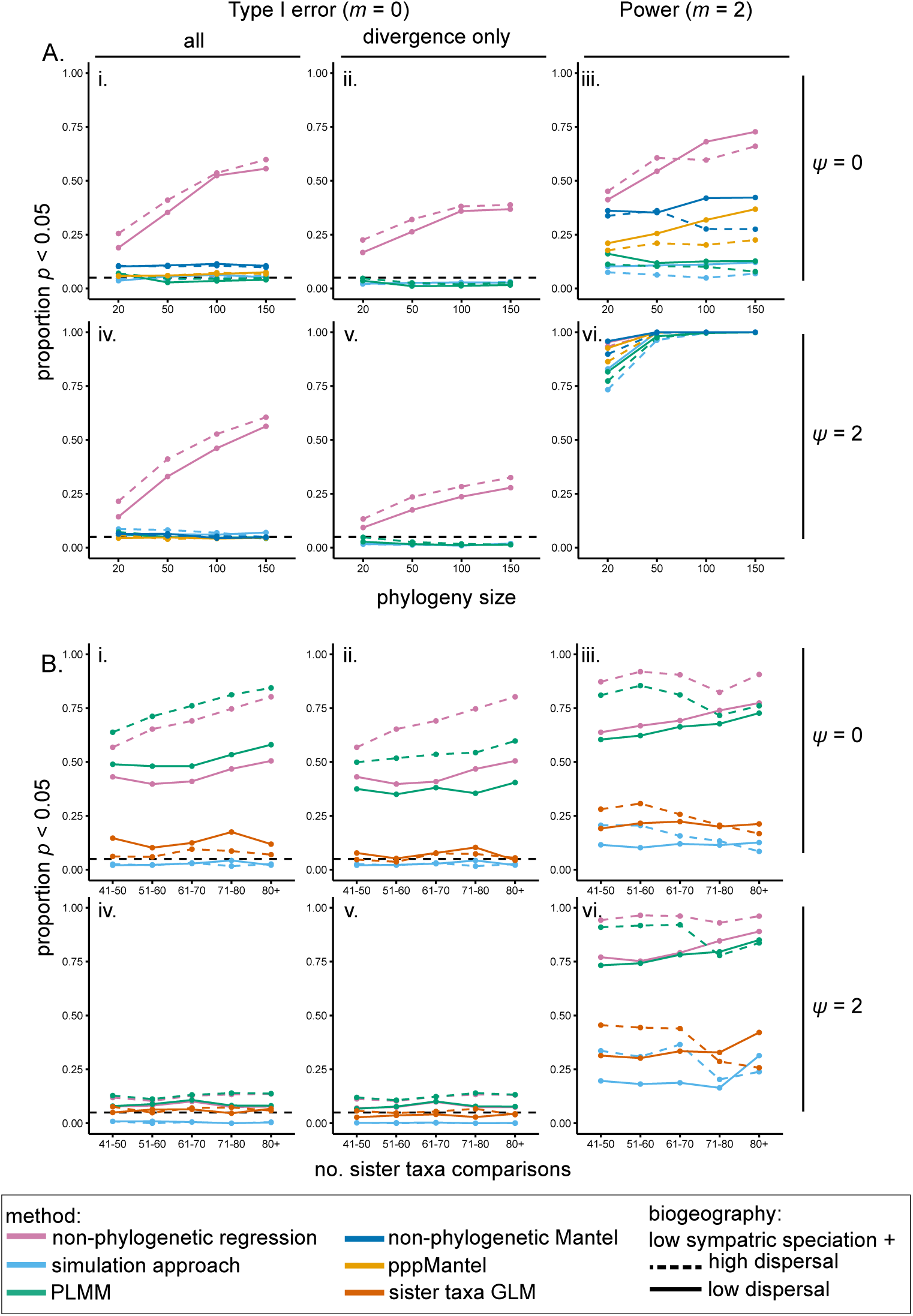
Proportion of statistically significant analyses in datasets simulated under divergent character displacement in biogeographic scenarios with low sympatric speciation rates. A. Results from approaches using data from all pairwise comparisons in a clade, plotted as a function of the phylogeny size and dispersal rate when i-ii. *m* = 0 and *ψ* = 0 (i. all analyses and ii. only analyses returning divergence in sympatry), iii. *m* = 2 and *ψ* = 0, iv-v. *m* = 0 and *ψ* = 2 (iv. all analyses and v. only analyses returning divergence in sympatry), and vi. *m* = 2 and *ψ* = 2. B. Results from analyses of sister-taxa culled from complete phylogenies binned by the number of resulting species pairs, plotted as a function of the number of sister taxa comparisons and dispersal rate when i-ii. *m* = 0 and *ψ* = 0 (i. all analyses and ii. only analyses returning divergence in sympatry), iii. *m* = 2 and *ψ* = 0, iv-v. *m* = 0 and *ψ* = 2 (iv. all analyses and v. only analyses returning divergence in sympatry), and vi. *m* = 2 and *ψ* = 2. For scenarios where *m* = 2, only the proportion of significant results showing divergence are plotted. Dashed horizontal lines represent a Type I error rate of 5%.

The ability to detect divergence was relatively similar for *m* = 1 and *m* = 2, but declined for *m* = 10 (Fig. S5). This is due to a positive relationship between the ability to detect character displacement and the ratio of *ψ:m* (Fig. S6), resulting from a higher absolute magnitude of repulsion when both processes are present (Figs. S4, S6), indicating that this ratio impacts the ability to detect divergence more than the raw value of *m*.

For sister-taxa analyses, there is a high probability of falsely concluding that character displacement occurred in datasets simulated under BM and, to a lesser extent, OU, whether analyzed with simple linear regressions, sister-taxa GLMs, or PLMMs (Figs. 2Bi,iii, S2Bi,iii). As with the whole-tree approach, the power tends to increase and Type I error rate tends to decrease in datasets with attraction toward a single-stationary peak (Figs. 2Biv-vi, S2Biv-vi). However, the overall power to infer the presence of divergence was low with sister-taxa analyses (Figs. 2Biii,vi, S2Biii,vi). Inferences were generally better when dispersal was high, which may reflect the elevated observed divergence in high dispersal scenarios (Fig. S3). Allopatric speciation scenarios increased the probability of Type I error (Fig. 2Bi-ii).

For the phylogenetic trait model-fitting analyses, BM and OU were generally correctly chosen when they were the generating models (i.e., when *m* = 0 and when *ψ* = 0 or 2, respectively, Figs. 3, S8). When *ψ* = 0 and *m* > 0, the matching competition (MC) model with biogeography is consistently the best-fit model (Figs. 3A, S8A). When *m* > 0 and *ψ* =2, the diversity dependent exponential (DD_exp_) model with biogeography was favored over other models in most scenarios (Figs. 3B, S8B), with positive rate parameters estimated in the maximum likelihood solution (Fig. S9). The biogeographic scenario did not greatly affect the outcome of model fitting, though correct models were slightly more supported when dispersal was high (Fig. S10), again in agreement with the observed magnitude of repulsion (Fig. S3). Although the models are less identifiable when *m* = 10 and *ψ* = 2 (Figs. 3, S8), this results from variation in the *ψ:m* ratio— there is a ratio of *ψ:m* around which these models cannot be distinguished (Fig. S11).

**Figure 3.**
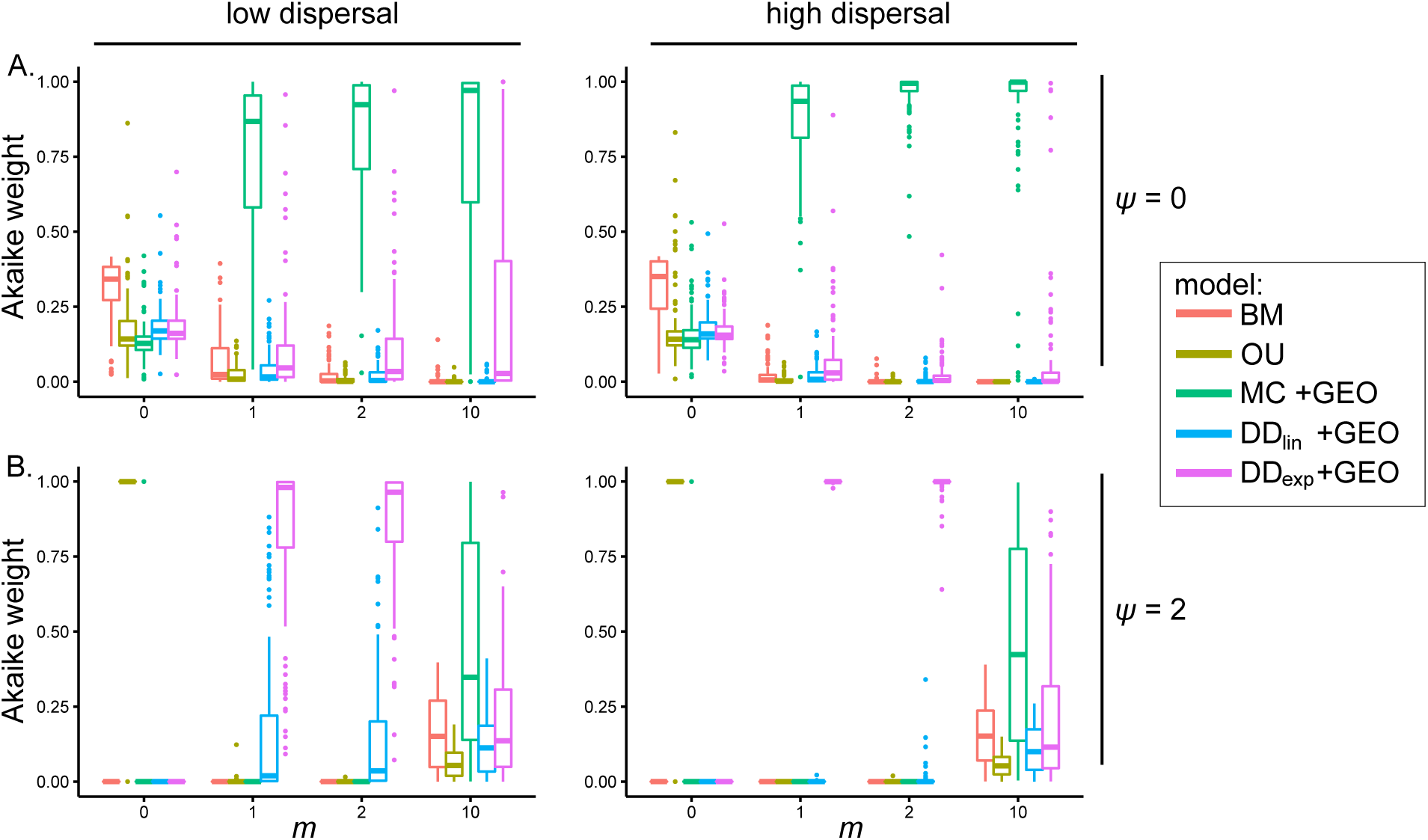
Boxplots of Akaike weights for each trait model fit to simulated datasets in biogeographic scenarios with low sympatric speciation rates as a function of *m* in trees with 100 species. A. When OU is absent, BM is the best-fit model when *m* = 0, and the matching competition model with biogeography is the best model when competitive divergence is present. B. When OU is present, OU is the best-fit model when *m* = 0, and the diversity-dependent exponential model with biogeography is the best model when competitive divergence is present and *ψ:m* is relatively high.

Process-based models fit to sister-taxa datasets in EvoRAG did not mistakenly identify an effect of species interactions when they were absent (Fig. S4A, C, Table 2), but they were unable to identify the effect of competition when *ψ* = 0 (Fig. S4B, Table 2). However, as with process-based models fit to the whole phylogeny, when data were simulated with both repulsion and a pull toward a stable peak, a model where evolutionary rates vary linearly with the number of sympatric taxa is often the best-fit model, though generally with only a marginally lower AICc value (i.e., ∆AICc < 2) than BM (Fig. S4, Table 2).

### Convergent Character Displacement

As with divergent character displacement, with all pairwise species combinations, the ability of most methods to detect convergent character displacement depends on the presence of the OU process on the resource-use trait: datasets simulated under an OU model were more likely to be statistically significant (Figs. 4A.iv-vi, S12A.iv-vi) across all methods than those with BM simulated resource-use traits (Figs. 4A.i-iii, S12A.i-iii). Again, this is likely because the presence of the OU process in the resource-use trait amplifies the magnitude of convergence (Fig. S13, S14). Overall, however, only the simulation approach had substantial power (> 0.80) to detect convergent character displacement (Table 1), and only in trees with 100 or more tips and datasets with the OU process in the simulated resource-use trait. Indeed, the nonphylogenetic regressions often (spuriously) detected divergence rather than the simulated convergence, especially in smaller trees (Supplementary Tables). Both types of Mantel tests were unable to detect convergence, in fact having a higher Type I error rate (detecting divergence in BM simulated datasets, Supplementary Tables) than power. As with divergent character displacement, there was a tendency for higher power in lower dispersal scenarios.

**Figure 4.**
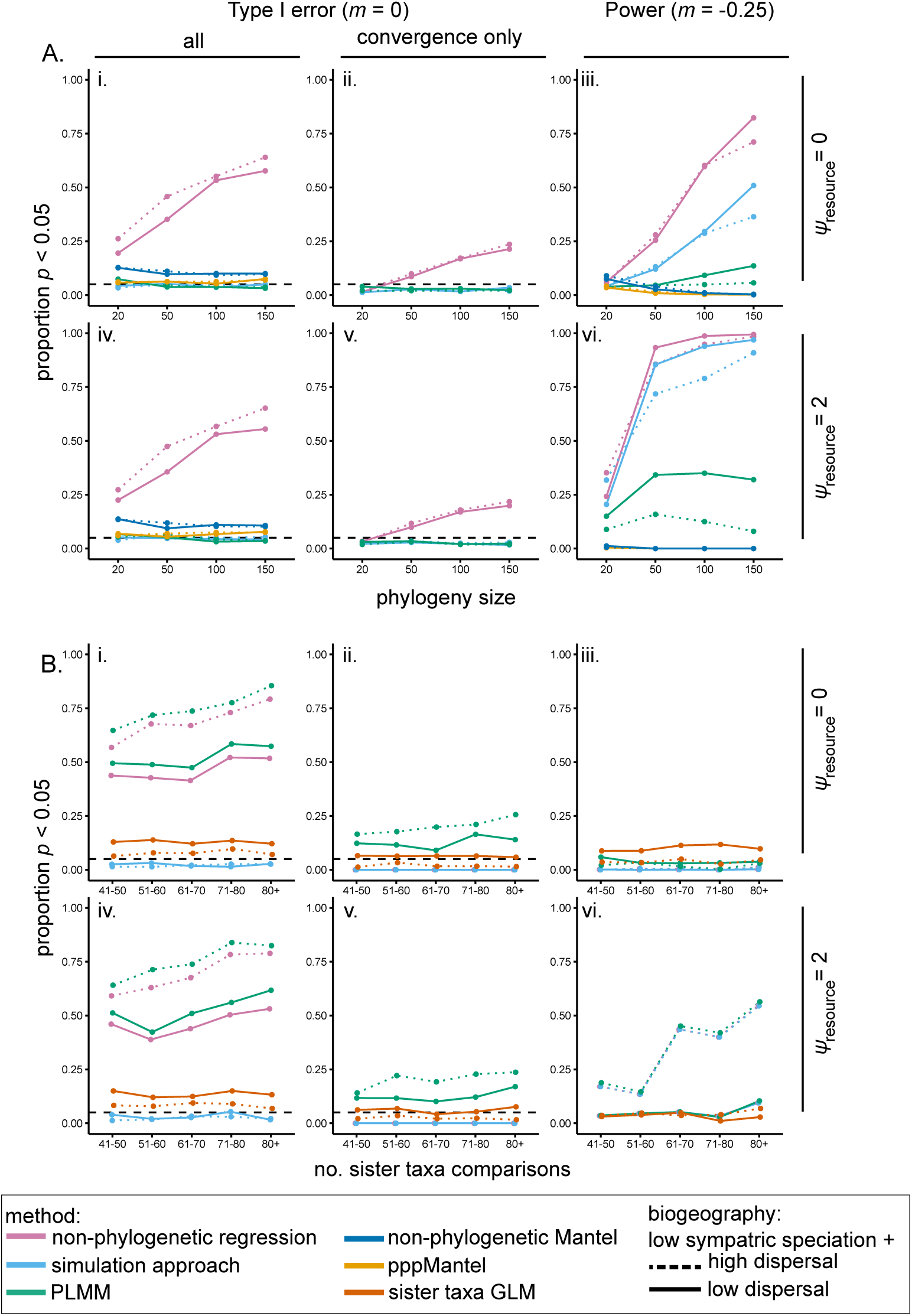
Proportion of statistically significant analyses in datasets simulated under convergent character displacement in biogeographic scenarios with low sympatric speciation rates. A. Results from approaches using data from all pairwise comparisons in a clade, plotted as a function of the phylogeny size and dispersal rate when i-ii. *m* = 0 and *ψ_resource_*= 0 (i. all analyses and ii. only analyses returning convergence in sympatry), iii. *m* = −0.25 and *ψ_resource_* = 0, iv-v. *m* = 0 and *ψ_resource_* = 2 (iv. all analyses and v. only analyses returning convergence in sympatry), and vi. *m* = −0.25 and *ψ_resource_* = 2. B. Results from analyses of sister-taxa culled from complete phylogenies binned by the number of resulting species pairs, plotted as a function of the number of sister taxa comparisons and dispersal rate when i-ii. *m* = 0 and *ψ_resource_* = 0 (i. all analyses and ii. only analyses returning convergence in sympatry), iii. *m* = −0.25 and *ψ_resource_* = 0, iv-v. *m* = 0 and *ψ_resource_* = 2 (iv. all analyses and v. only analyses returning convergence in sympatry), and vi. *m* = −0.25 and *ψ_resource_* = 2. For scenarios where *m* = −0.25, only the proportion of significant results showing convergence are plotted. Dashed horizontal lines represent a Type I error rate of 5%.

**Table 1.** Summary of the statistical properties of the analytical approaches tested under scenarios using data from all tips (i.e., with sister-taxa analyses excluded). Values refer to the range of type I error rates and power levels for each tree size ≥50, averaged across biogeographic scenarios and scenarios where *ψ* or *ψ_ηf_* = 0 or 2. Power refers to only those statistically significant tests in the appropriate tail (i.e., in the lower tail for divergent character displacement and upper tail for convergent character displacement). For each analytical scenario, the cell with the method with the best trade-off between Type I error and power is highlighted.

| Analysis | non-phylogenetic regression |  | Mantel test |  | pppMantel test |  | PLMM |  | simulation test |  | process-based models |  |
| --- | --- | --- | --- | --- | --- | --- | --- | --- | --- | --- | --- | --- |
|  | *type I* | *power* | *type I* | *power* | *type I* | *power* | *type I* | *power* | *type I* | *power* | *type I* * | *power*<sup>†</sup> |
| divergent char. displacement | 0.37-0.61 | 0.51-1 | 0.05-0.10 | 0.29-1 | 0.04-0.06 | 0.19-1 | 0.05-0.06 | 0.12-1 | 0.05-0.07 | 0.08-1 | 0.04-0.05 | 0.91-0.94 |
| convergent char. displacement | 0.40-0.60 | 0.31-0.99 | 0.08-0.09 | 0-0.02 | 0.05-0.06 | 0-0.01 | 0.05-0.07 | 0.07-0.26 | 0.04-0.05 | 0.12-0.91 | -- | -- |
| predicting spp. interactions | 0.08-0.3 | 1 | -- | -- | -- | -- | 0.07-18 | 1 | 0.03-0.04 | 1 | -- | -- |

\*Type I error rate calculated as the proportion of datasets simulated without divergence for which a model that includes species interaction— DD<sub>exp</sub>, DD<sub>lin</sub>, or MC—was chosen by model selection (i.e., for which $\Delta AICc = 0$ and $\Delta AICc$ for all other models $> 2$ ).
<sup>†</sup> Power calculated as the proportion of datasets simulated with divergence for which either DD<sub>exp</sub>, DD<sub>lin</sub>, or MC was chosen by model selection.

The power to detect convergence generally increased with increasingly negative values of m, the maximum strength of attraction in the signal trait when species are identical in the resource-use trait (Fig. S15), though as *m* gets large, the probability that all species converge on the same trait value increases, especially when *ψ*_resource_ = 2 (data not shown).

Regardless of whether resource-use traits are simulated under OU or BM, when there is no convergence, most methods used for sister-taxa analyses tend to have high Type I error rates, though these analyses return an erroneous inference of divergence, rather than convergence, between sister taxa (Figs. 4B.i,ii,iv,v, S12B.i,ii,iv,v, Supplementary Tables). Sister-taxa analyses had overall low power to detect convergence when it did exist, and non-phylogenetic regressions often detected divergence, rather than convergence (Supplementary Tables). When convergence was detected, it tended to be in biogeographic scenarios with high dispersal, likely reflecting the overall magnitude of convergence achieved (Fig. S13). As with divergent character displacement simulations, the allopatric speciation biogeographic scenarios were more likely to lead to higher Type I error rates (Figs. 4B.i,iv). Process-based models fit to sister-taxa datasets in EvoRAG did not erroneously detect divergence or convergence (i.e., BM was the best-fit model when *m* = 0, Fig. S14 A, C, Table 2), but they could not detect an effect of species interactions when convergence was present, at least for the number of sister taxa in this study, as OU was the best-fit model when *m* = −0.25 (Fig. S14 B, C, Table 2).

**Table 2.**
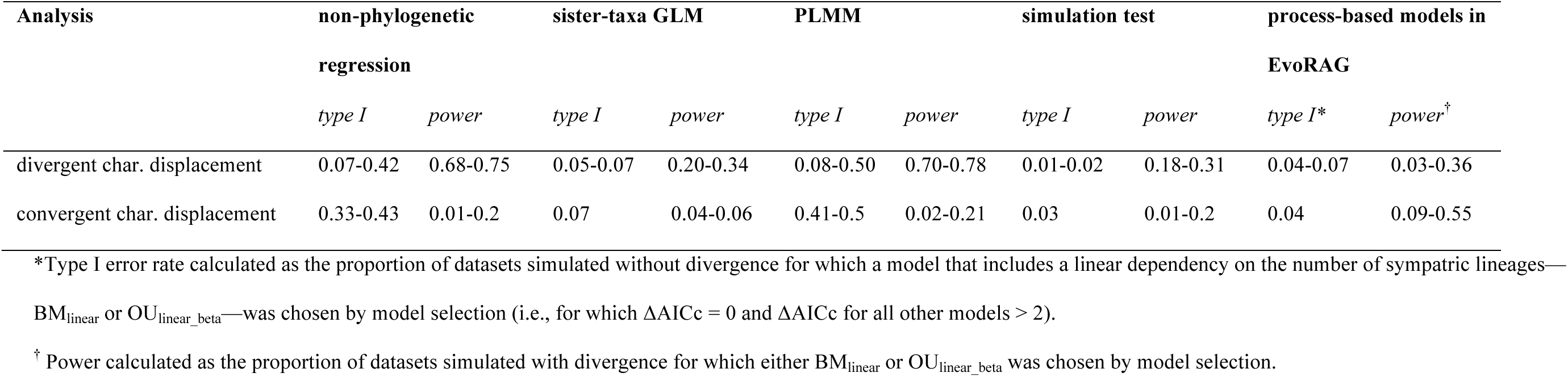
Summary of the statistical properties of the analytical approaches tested under scenarios using sister-taxa analyses. Values refer to the range of type I error rates and power levels, averaged across biogeographic scenarios and scenarios where *ψ* or *ψ_nf_* = 0 or 2. Power refers to only those statistically significant tests in the appropriate tail (i.e., in the upper tail for divergent character displacement and lower tail for convergent character displacement). We caution against using sister-taxa approaches to test for character displacement.

| Analysis | non-phylogenetic regression |  | sister-taxa GLM |  | PLMM |  | simulation test |  | process-based models in EvoRAG |  |
| --- | --- | --- | --- | --- | --- | --- | --- | --- | --- | --- |
|  | *type I* | *power* | *type I* | *power* | *type I* | *power* | *type I* | *power* | *type I* * | *power*<sup>†</sup> |
| divergent char. displacement | 0.07-0.42 | 0.68-0.75 | 0.05-0.07 | 0.20-0.34 | 0.08-0.50 | 0.70-0.78 | 0.01-0.02 | 0.18-0.31 | 0.04-0.07 | 0.03-0.36 |
| convergent char. displacement | 0.33-0.43 | 0.01-0.2 | 0.07 | 0.04-0.06 | 0.41-0.5 | 0.02-0.21 | 0.03 | 0.01-0.2 | 0.04 | 0.09-0.55 |

\*Type I error rate calculated as the proportion of datasets simulated without divergence for which a model that includes a linear dependency on the number of sympatric lineages— $BM_{linear}$ or $OU_{linear\_beta}$ —was chosen by model selection (i.e., for which $\Delta AICc = 0$ and $\Delta AICc$ for all other models $> 2$ ).
<sup>†</sup> Power calculated as the proportion of datasets simulated with divergence for which either $BM_{linear}$ or $OU_{linear\_beta}$ was chosen by model selection.

### Predicting Interspecific Interactions

Although all three methods used to identify traits that are causally related to interspecific interactions had high power (>>0.8, Table 1, Supplementary Tables) to do so in the parameter space explored here (Figs. 5ii,iv, S16ii,iv), only the simulation approach had both high power and a low Type I error rate (Table 1), whereas non-phylogenetic regressions and PLMMs had fairly high Type I error rates (Table 1) when interactions were simulated based on similarity in a trait other than the measured one (Fig. 5i,iii, S16i,iii). The power to detect an interaction was not greatly affected by the coefficients used to simulate datasets (Fig. S17). Biogeography did not have a large impact on analyses, though there were slightly higher Type I error in low-dispersal scenarios (Fig. 5i,iii).

**Figure 5.**
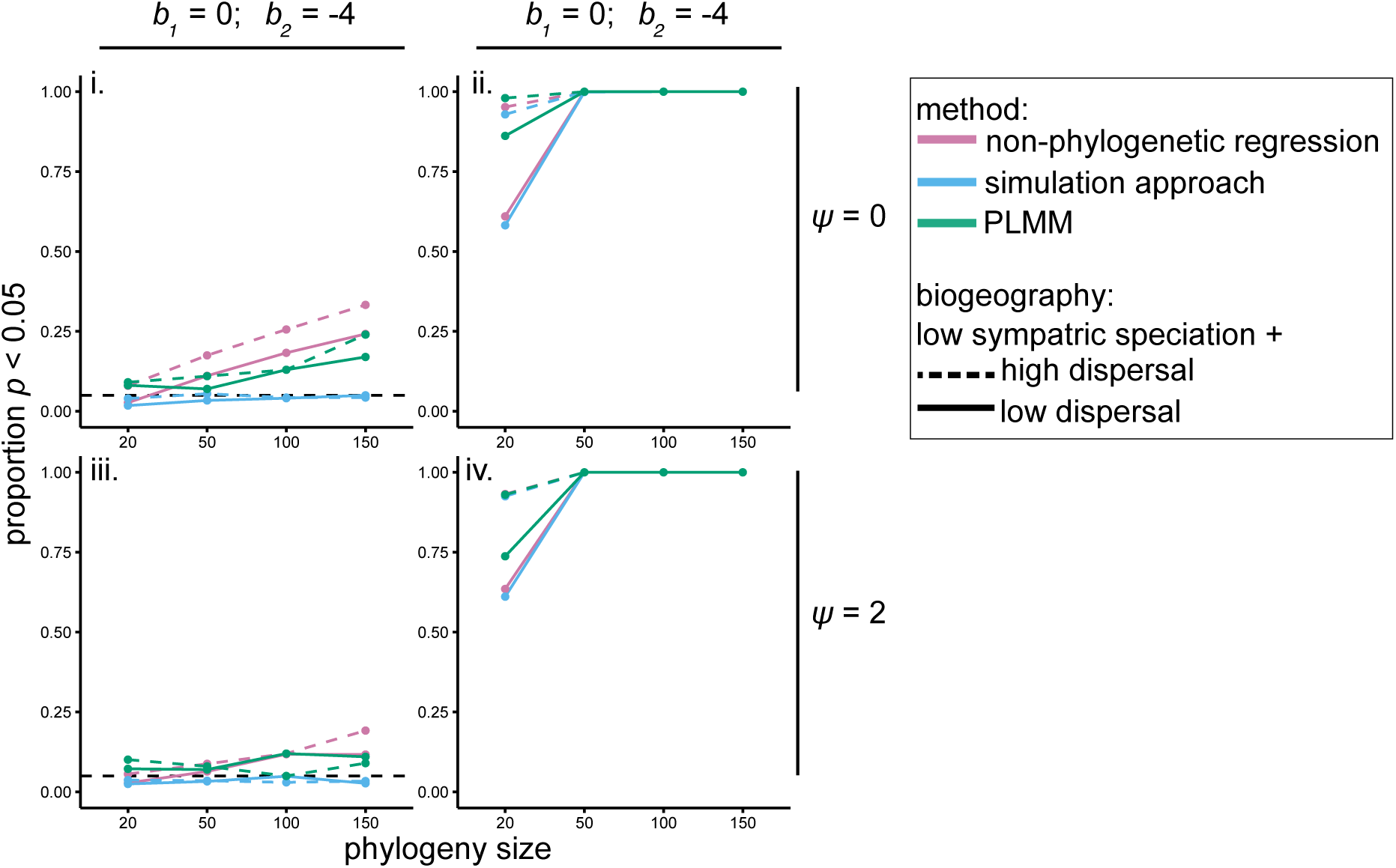
Proportion of statistically significant analyses in datasets with interactions simulated under a simple phenotype matching process in biogeographic scenarios with low sympatric speciation rates. Results from analyses where the measured trait was simulated under BM (i, ii) or OU (iii, iv), plotted as a function of the phylogeny size and dispersal rate when i. *b_1_* (the simulation coefficient determining the relationship between the interaction and the measured trait) = 0, *b_2_* (the simulation coefficient for an unmeasured trait) = −4, and *ψ* = 2, ii. *b_1_* = −4, *b_2_* = 0, and *ψ* = 2, iii. *b_1_* = 0, *b_2_* = −4, and *ψ* = 0, and iv. *b_1_* = −4, *b_2_* = 0, and *ψ* = 0.

## DISCUSSION

As open-access databases with species range, trait, and phylogenetic data rapidly expand, investigators are able to test hypotheses about the relationships between interspecific interactions and phenotypic evolution at an unprecedented scale. Understanding the relative strengths and weaknesses of phylogenetic comparative methods available for testing such hypotheses is thus paramount. We found that currently used methods for detecting causal relationships between interspecific interactions and species phenotypes suffer from severe limitations (Tables 1,2).

Overall, standard methods are better at detecting divergent character displacement when divergence does not drive unbounded trait evolution (i.e., when selection acts against extreme phenotypes, as can be modeled by the OU process). Consistent with previous reports (Harmon and Glor 2010; Guillot and Rousset 2013), Mantel tests had high Type I error rates and both standard and pppMantel tests have low power (Table 1, Figs. 2Ai, S2Ai). Alarmingly, we found that commonly used sister taxa approaches have high Type I error rates (Table 2, Figs. 2Bi,iv, S2Bi,iv, 4Bi,iv, S12Bi,iv, Supplementary Tables), which would lead investigators to conclude that divergent character displacement had occurred when, in fact, it had not, and none have a reasonable combination of Type I error and power. Thus, we discourage empiricists from using sister-taxa approaches to study character displacement. If no other data are available for testing for character displacement on the whole tree, then we recommend phylogenetic simulations, as they are the only method with reasonably low type I error rates, even though they suffer from low power (Tables 1, 2).

Fitting process-based phylogenetic trait models to datasets simulated with divergent character displacement yielded more consistent patterns (Fig. 3). Without attraction toward a single stationary peak to bound trait evolution, the matching competition (MC) model with biogeography was predominantly the best-fit model. For datasets simulated with the OU process, the diversity-dependent exponential (DD_exp_, see Box 1) model with biogeography was the best-fit model, and similarly a model with a linear relationship between evolutionary rates and the number of sympatric taxa often fit sister-taxa datasets, though with much lower power overall (Fig S4, Table 2). In the DD_exp_ model, rates of trait evolution vary exponentially with the number of sympatric lineages through time, so incorporating the effect of interspecific interactions on the rate of trait of evolution but not explicitly modeling the process of character displacement acting on the mean trait values. It may nonetheless provide a useful proxy for detecting patterns that are similar to those left by character displacement, in the absence of a process-based model that incorporates both attraction toward an optimum trait value and divergent character displacement. We emphasize, however, that statistical support for the DD_exp_ model does not in itself constitute decisive evidence that character displacement has occurred, as other processes may generate increasing evolutionary rates with increasing lineage diversity. Given that the DD_exp_ model is the best-fit model in parameter space where other methods also perform well, combined evidence from model-fitting and other, non-process based methods would constitute a strong case for the presence of character displacement. In the absence of tip data (e.g., due to incomplete sampling or traits that are inherently measured as pairwise properties), process-based models are unsuitable and we recommend using data from as many species pairs as possible—not just sister taxa—and using simulation approaches or PLMMs. In other words, to detect divergent character displacement, we recommend that empiricists fit the MC model to their dataset when possible. High support for the MC model would constitute evidence that character displacement has acted on a trait. If the MC model does not provide a good fit for the data, this could be because character displacement proceeds in the presence of bounded trait evolution, in which case a signature of the DD_exp_ model with a positive rate parameter and/or a signature of sympatric divergence in phylogenetic simulations or PLMMs would constitute evidence consistent with divergent character displacement.

Interestingly, even though most previous investigators have used the DD_exp_ model to represent a decline in ecological opportunity with increasing species richness (Mahler et al. 2010; Weir and Mursleen 2013), the maximum likelihood estimates of the rate parameters for this model were positive, rather than negative, when both divergence and the OU process were present (Fig. S9). This is consistent with our finding of increasing evolutionary rates with increasing species richness (Figs. S3, S4, S7) in this scenario. An increase in the rate of evolutionary changes in trait values toward the present likely results from selection not only restricting species to certain trait space but also partitioning that space. The resulting adaptive landscape is therefore rapidly changing, causing accelerating evolutionary rates as lineages fill this increasingly constrained space.

The MC model (Box 1) is similar to the model used to simulate data (Eq. 1), with the assumption that *α* is very small (<< 1) and consequently, competitive interactions are affected by the mean trait values of all sympatric species, rather than by pairwise similarity (Nuismer and Harmon 2015; Drury et al. 2016). Biologists, however, generally assume that competition is stronger between phenotypically similar species (Brown and Wilson 1956). Our results show that the assumption of a small *α* does not render the MC model useless for studying character displacement, as the MC model is the best-fit model for many datasets simulated under the character displacement model used here. Nevertheless, the finding that the DD_exp_ model is the best-fit model in datasets simulated under character displacement including OU indicates that the MC model is not a perfect model of character displacement. Recently, approximate Bayesian computational (ABC) tools have been published to fit a model of character displacement in which, like in our simulation model, the strength of competition depends on similarity in trait space (Clarke et al. 2017). This model provides an alternative tool for detecting character displacement in comparative datasets, and we hope that further development of methods such as this ABC method will help ameliorate the statistical issues shown here.

For datasets simulated including the OU process, the ratio of the pull-parameter in the OU portion of the model to the maximum amount of repulsion (*ψ:m)* had a consistent impact across all methods, which results from the overall larger magnitude of evolutionary changes in traits in scenarios with a high *ψ:m* ratio (Figs. S3, S4, S7). As *ψ:m* approached 1, all methods were better at detecting character displacement. Currently, there are no analytical approaches that can disentangle the simultaneous impact of attraction toward a peak and divergence due to competition, though we hope our results will inspire development of such tools.

Unlike for divergent character displacement, available statistical methods for detecting convergence in comparative datasets generally do a poor job of detecting convergence, with the simulation method outperforming others (Table 1). With whole-dataset approaches, Type I error rates are acceptable for phylogenetic analyses (~5%), however, so although detecting convergence is difficult, the risk of mistakenly detecting convergence is low. In sister-taxa analyses, although Type I error rates are high for PLMMs (Table 2), these largely return erroneous divergence results, rather than erroneous convergence (Figs. 4Bii,v, S12Bii,v). In short, if an empiricist detects convergence in their dataset, they can be fairly confident in the result. Yet if empiricists do not detect convergence, this could simply be a result of lower power of the available analytical tools. Currently, there are no tools to fit phylogenetic trait models of convergence between species (e.g., Nuismer & Harmon 2015); such tools might more successfully identify convergent character displacement in comparative datasets than the available statistical methods.

For both divergent and convergent character displacement scenarios, we found that sister-taxa GLMs and the simulation approach had a mean Type I error rate near 5% (Table 2). However, in some scenarios, the Type I error for sister taxa GLMs was much higher than for the simulation approach (Figs. 2Bi, 4Bi, Supplementary Tables), which suggests that including a model-based estimate of the rate of trait evolution more properly accounts for the effect of divergence than simply including the branch lengths separating sister taxa as a covariate in analyses to control for variation in the amount of time sister taxa have had to diverge from one another. The high overall Type I error rate for sister-taxa analyses may also result from the unrealistic assumption that transitions between allopatry and sympatry are uncommon along branches connecting sister taxa (Weir and Price 2011; Tobias et al. 2014). Supporting this explanation, we found that biogeographic scenarios with high levels of sympatric speciation and low dispersal tended to have overall lower Type I error rates (cf. Figs. 2,S2; Figs. 4,S12).

The outlook for identifying which traits drive species interaction is brighter. The statistical methods available to test for causal relationships between phenotypic similarity and interactions between species have very high power. The simulation approach has a low Type I error rate when causal relationships are simulated based on an unmeasured trait, although nonphylogenetic regressions and PLMMs suffer from relatively high Type I error rates (Table 1). Thus, we recommend that empiricists interested in predicting pairwise species interactions based on trait data use phylogenetic simulations. While we did not simulate interactions between clades, our results are likely applicable to other empirical questions, such as identifying traits that predict links in ecological networks (Rafferty and Ives 2013; Hadfield et al. 2014; Eklöf and Stouffer 2016).

By simulating datasets with various types of interactions between species across different modes of speciation and dispersal rates, we have shown that many of the methods that investigators use to analyze empirical datasets have low power to detect such patterns (Table 1). Worse still, widely-used sister taxa approaches, including standard regressions and sister-taxa GLMs, often detected character displacement in datasets that were simulated under a simple BM model (Figs. 2Bi-ii, 4Bi-ii). We therefore urge investigators to use caution when interpreting the results of such analyses, even in cases when sympatry is delineated using other criteria than the one considered here. When process-based models could be fit to these datasets, they tended to consistently identify patterns of divergence (i.e., either the matching competition model or a diversity-dependent model is the best fit model >91% of the time). Thus, when possible, empiricists should employ such methods. Statistical tools to fit process-based models of phenotypic evolution including species interactions are in their infancy (Drury et al. 2016; Manceau et al. 2017) and many possible models are not yet available (e.g., convergent character displacement, character divergence in the presence of an adaptive pull towards a peak). We hope that our results encourage the continued development of such tools.

In closing, we note that divergent character displacement is erroneously detected with many statistical approaches, indicating that there may be an overrepresentation of empirical studies that imply that divergence has occurred. In particular, studies that have used sister-taxa methods to document character displacement may have falsely interpreted a null expectation— larger trait differences between sympatric lineages owing to allopatric speciation—as evidence for divergent character displacement. Conversely, convergent character displacement is often hard to detect with existing methods, suggesting that convergence in signal traits (e.g., Cody 1969, 1973; Tobias et al. 2014; Losin et al. 2016) might be more prevalent than previously thought.

## ACKNOWLEDGEMENTS

This research was funded by the European Research Council (grant 616419-PANDA to HM) and the National Science Foundation (grant DEB-1457844 to GFG). We thank M. Manceau, J. Clavel, E. Lewitus, O. Maliet, and O. Missa for feedback and F. Hartig for assistance streamlining our simulation script.

## Box 1. Methods for assessing the interplay between interspecific interactions and species phenotypes

Comparative analyses of the interplay between interspecific interactions and species phenotypes can either be conducted on entire clades, or, commonly, on sister taxa—species pairs that share a most recent common ancestor—that are culled from larger phylogenies. Such analyses generally consist of testing the statistical significance of correlations between either phenotypic similarity and geographic overlap (to test for divergent or convergent character displacement) or species interactions and phenotypic similarity (to find predictors of species interactions). As we are looking for correlations between pairwise comparisons (e.g., trait similarity, biogeographical overlap, hybridization, magnitude of pre-zygotic isolation), rather than “tip values” belonging to a single species, phylogenetically independent contrasts and extensions of PGLS analyses (Felsenstein 1985; Rezende and Diniz-Filho 2012) cannot be used, and alternative tests have been developed.

### 1. Non-phylogenetic regressions

“Non-phylogenetic regressions” refers to Generalized Linear Models (GLMs) that ignore phylogenetic structure. Though less commonly applied to whole-clade analyses, investigators sometimes use non-phylogenetic regressions for sister-taxa analyses, on the basis that branches connecting sister taxa represent independent evolutionary histories (Felsenstein 1985).

### 2. Mantel tests

Several previous investigators have implemented Mantel tests (Mantel 1967) in analyses of species-pair comparisons (e.g., Roncal *et al.* 2012). These tests are designed to assess correlations between matrices, which here comprise interspecific trait distances or differences. Existing accounts of Mantel tests describe procedures only for complete matrices, so they cannot be used in many cases, including sister-taxa analyses (for which most off-diagonal elements of distances matrices are by definition excluded) and in identifying predictors of species interactions (e.g., hybridization), as only sympatric lineages can interact and setting values for allopatric comparisons to zero would not make biological sense.

### 3. Phylogenetically permuted partial Mantel tests

Phylogenetically permuted partial Mantel (pppMantel) tests (Lapointe and Garland 2001) account for phylogenetic non-independence by permuting null datasets that are structured phylogenetically, and are popular among investigators studying species interactions (e.g., Allen *et al.* 2014; Willis *et al.* 2014; Medina-García *et al.* 2015). Like Mantel tests, pppMantel tests also require complete interaction matrices.

### 4. Phylogenetic linear mixed models

In recent years, researchers have adapted animal models from quantitative genetics to incorporate phylogenies as random effects in mixed-effect regressions on comparative datasets (Hadfield & Nakagawa 2010). Such phylogenetic linear mixed models (PLMMs) have been modified to accommodate pairwise species data (Tobias et al. 2014), wherein the identity of the species being compared and the node connecting them in the phylogeny are included as random effects. PLMMs are promising new tools, as they are not limited to sister-taxa data and model predictions can be generated and plotted.

### 5. Phylogenetic simulations

Simulation approaches are widely used to control for phylogenetic non-independence in tip data (Martins & Garland Jr 1991; Garland *et al.* 1993), and have been applied to pairwise species comparisons (Elias et al. 2008; Drury et al. 2015; Losin et al. 2016). In these approaches, trait evolution is simulated along phylogenies, often scaled such that the simulated tip data resemble real data. Pairwise comparisons are then calculated on many simulated datasets and used to generate a phylogenetically informed null distribution of test statistics against which to compare test statistics calculated from the real data.

### 6. Process-based models of phenotypic evolution

In the statistical approaches outlined thus far, the data analyzed are measurements of pairwise differences between species, and the statistical tests for the effect of species interactions on trait evolution consist of testing for significant correlations between either phenotypic similarity and geographic overlap or species interactions and trait similarity. However, it is also possible to detect a signature of interspecific competition in the distributions of continuous trait values across the tips of a phylogeny by fitting process-based models of phenotypic evolution to the data. These models allow testing hypotheses about which processes are most likely to have generated the observed distribution of traits in a clade (Hansen & Martins 1996).

Interspecific interactions have recently been incorporated into such models in two ways. First, in diversity-dependent (DD) models, evolutionary rates change as a function (either linear [DD_lin_] or exponential [DD_exp_]) of the number of extant lineages through time (e.g., Weir & Mursleen 2013). Secondly, in the ‘matching competition’ (MC) model, trait evolution in an evolving lineage varies as a function of the values of traits in other evolving lineages (Nuismer & Harmon 2015, Drury *et al.* 2016). Comparing the fit of these models to other models that exclude interspecific interactions (e.g., Brownian motion and Ornstein-Uhlenbeck models) tests whether there is evidence that interspecific interactions have influenced the trajectory of trait evolution in a clade.

### 7. Sister-taxa GLMs

If allopatric speciation is common, then sympatry occurs after a period of initial isolation, resulting in a pattern where sympatric sister taxa are older than allopatric sister taxa. Thus, even random genetic drift can generate a pattern in which sympatric lineages have more divergent traits compared to allopatric lineages, simply because divergence has had more time to evolve (Weir and Price 2011; Tobias et al. 2014). To control for variation in the evolutionary distance between sister taxa, “sister taxa GLMs” include patristic distance as a predictor in non phylogenetic regressions (e.g.,Davies *et al.* 2007; Martin *et al.* 2010).

### 8. Sister-taxa model fitting

Recently, tools have been described for fitting process-based models to sister taxa datasets using maximum likelihood (Weir and Wheatcroft 2011; Weir and Lawson 2015). With these tools, it is possible to test whether models that allow evolutionary rates to vary as a linear function of a gradient (e.g., whether male plumage coloration varies as a function of the strength of sexual selection, Seddon et al. 2013) better fit sister-taxa datasets than constant rates models. When the gradient is the number of sympatric lineages, these models are conceptually similar to the linear diversity dependent models described above.

## SUPPLEMENTARY METHODS

### Simulating Phylogenies Under Varying Biogeographic Scenarios

Joint simulation of trees and biogeographies requires parameterizing (1) the diversification process (the rules defining how new species appear), (2) rates of anagenetic range gains/losses (the rules defining how lineages’ ranges change), and (3) cladogenetic range inheritance (the rules defining how two sister lineages divide their ancestral range upon speciation). In each scenario, we evolved species ranges under the DEC+J model (Matzke 2014) across a grid of ten possible regions with equal probability of transitions between regions and only one region occupied at the root of the tree. We simulated the diversification process along each tree with a rate of 0.25 (speciation/lineage/time unit). Anagenetic changes were simulated under two dispersal rates {d (transition/lineage/unit time) = 0.03, 0.06}, chosen from within a range of dispersal values estimated from empirical datasets. When anagenetic dispersal happens, a lineage occupies an additional region chosen at random. Across these two values of “d”, we held local extirpation rates constant at e = 0.03 (local extirpation/lineage/unit time), the median value used for simulations in Matzke 2014. In cases where a lineage occupying only one region goes locally extinct, that lineage goes extinct (Matzke 2014).

The cladogenetic events possible in the DEC+J model are—briefly (for a detailed explanation of each of these processes, see Matzke 2014)— sympatric speciation (both daughter lineages keep the one-region ancestral range, parameter “y”), subset sympatric speciation (one of the daughter lineages keeps the ancestral range, the other one inherits a subset of the ancestral range, parameter “s”), vicariance (the two daughter lineages split the ancestral range, parameter “v”), and founder event speciation (one daughter lineages keeps the ancestral range, the other occupies a new region, parameter “j”).

As the pool of possible daughter ranges changes depending on the ancestral range (e.g., there are fewer ways in which the ancestral two-region range “AB” can be propagated to daughter lineages than the three-range region “ABC”), cladogenetic range changes are sampled by first assigning each type of cladogenetic change (y,s,v, and j from above) a particular weight. From these weighted ranges, the probability that a daughter inherits a specific range is calculated by dividing the weight assigned to that range by the sum of the weights of each possible daughter range (Matzke 2014). By default, the relative weight of sympatric speciation (range copying), subset sympatric speciation, and vicariance are equal in most implementations of the DEC model (i.e., y=s=v). To generate biogeographic scenarios where recently diverged sister taxa are more likely to be allopatric, we implemented a scenario where the weight of either type of sympatric speciation event is very low {y = s = 0.005*v} as well as under default parameter values {y = s = v}. Across both scenarios, we held the relative weight of founder event speciation constant (j = 0.1. y + s + v = 2.9).

### Simulating Character Displacement

Datasets were simulated under Eq. 1 in the main text as follows:

1. At the first time step (the root), the trait value was set to z_0_ (= 0 in all cases).
2. Each time step *dt* was set to the total tree height divided by 2500. To complete the simulation along any branch not divisible by this value of *dt,* we set the time step equal to the remainder of the branch length divided by *dt*.
3. During each time step from the root to the tip of the tree, the trait value of lineage *i* is calculated according to Eq 1. For the component of Eq.1 dictating the magnitude of divergence or convergence, between-lineage distances in trait space were calculated based on similarity at time *t-dt.* In simulations of divergent character displacement, if species have identical trait values, the “sign” value is overridden so that species move in the opposite direction of one another in trait space.
4. At a branching event, the trait values of both daughter lineages are set to equal the value of the parent lineage.

For divergent character displacement scenarios, we arbitrarily chose simulation parameter values based on visual inspection of simulated trajectories under different combinations of parameter values. We chose to focus on *m* = 2, and also explored the effect of varying this parameter between 1 & 10, because we could visualize divergence in the realized simulations. For example, for a 20 tip tree simulated with high dispersal and low sympatric speciation:

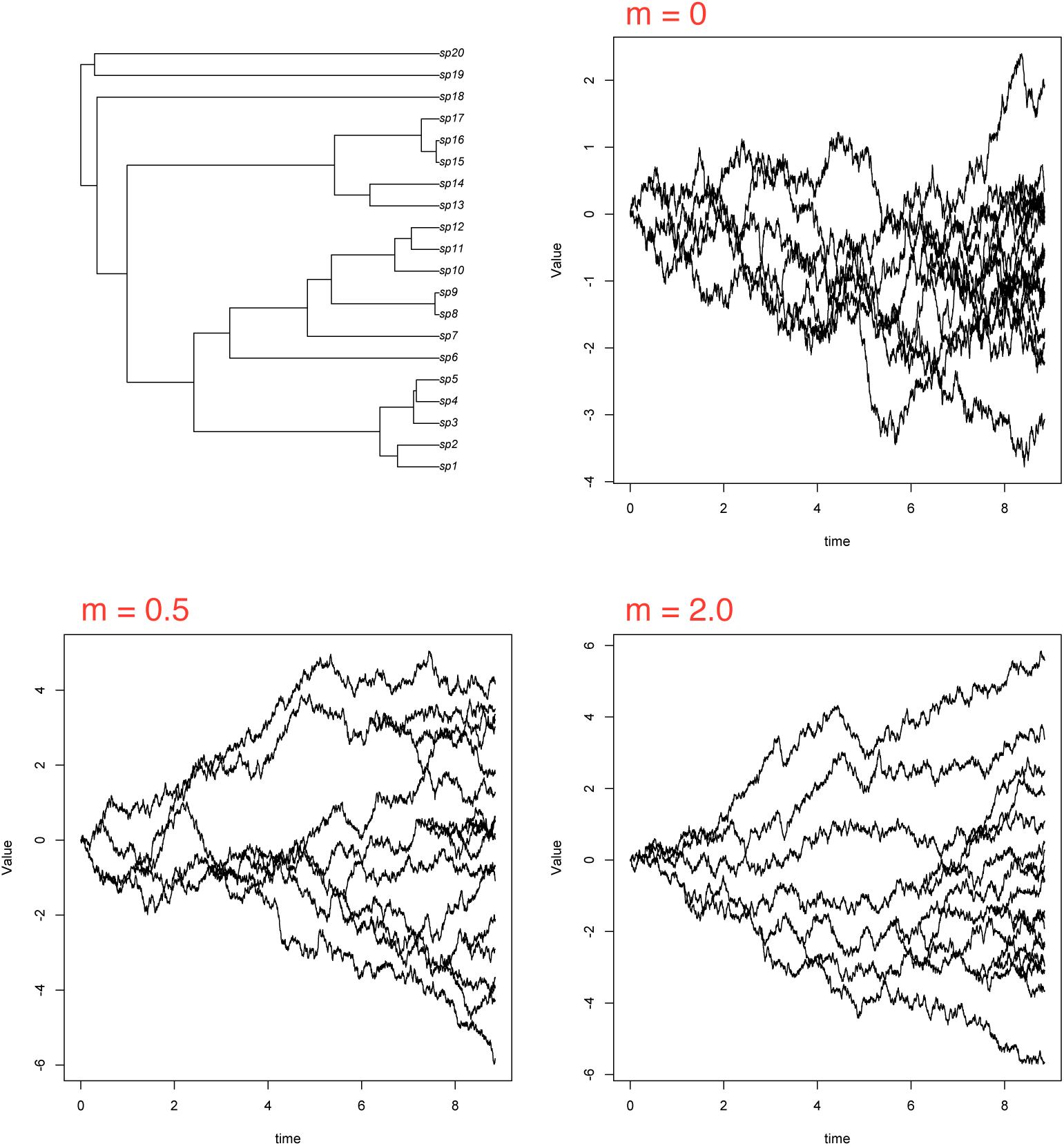

In convergent character displacement simulations, we likewise chose parameter values based on visual inspection of simulated trajectories under different combinations of parameter values. We chose to focus on *m* = −0.25, and also explored the effect of varying this parameter between −0.1 & −0.50, because we could visualize convergence in the realized simulations and the simulated trait values did not converge across the entire clade. For example, on the same tree as above:

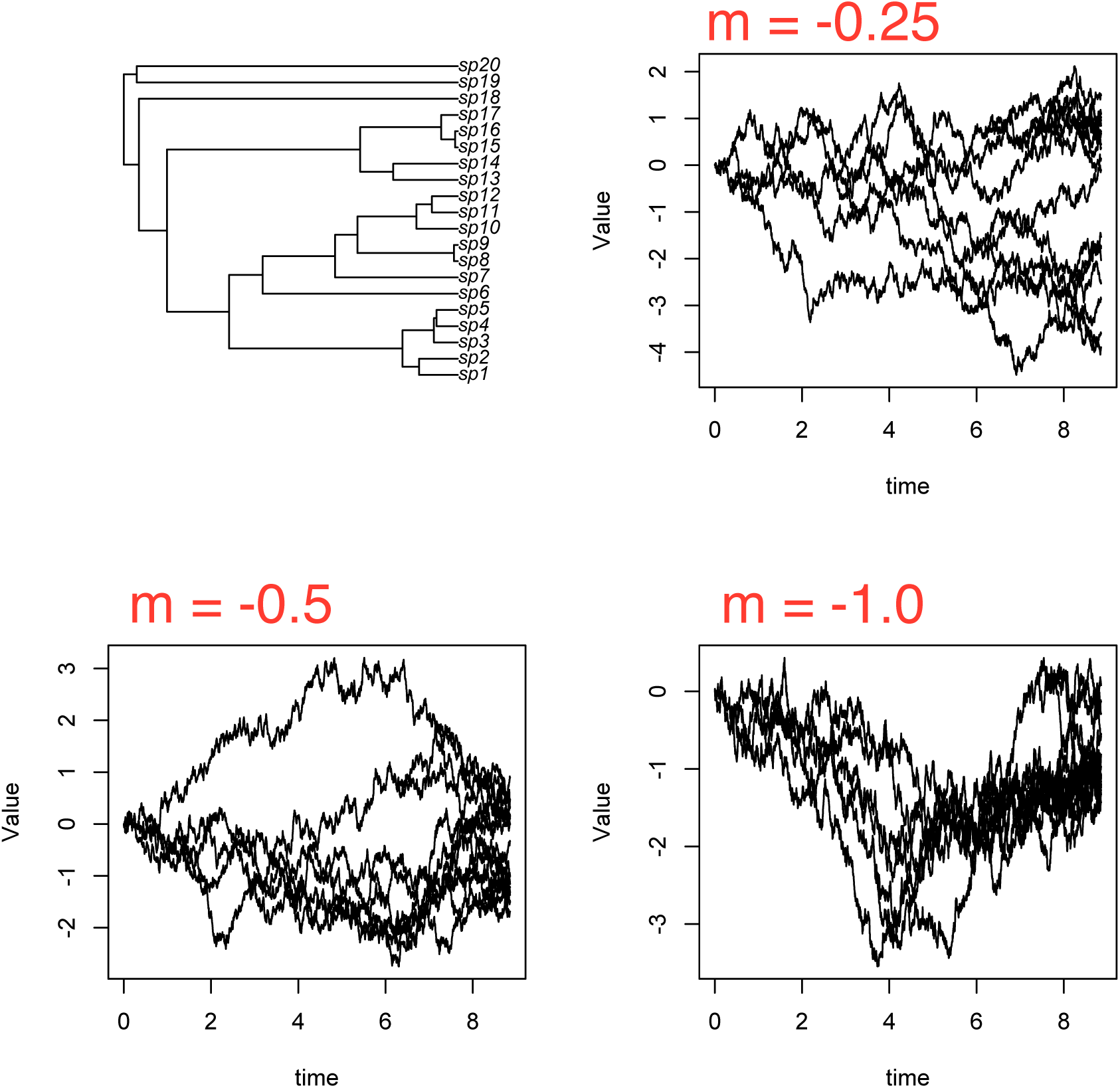

### Analytical Methods

Non-phylogenetic regressions: General Linear Models (GLMs) were fitted to simulated datasets using the glm function in R. For character displacement analyses, linear models were fit to pairwise differences in trait data, with sympatry/allopatry as the predictor variable. For interaction analyses, logistic regressions were fit to the simulated species interaction variable, with pairwise difference in the focal trait as the predictor variable.

Mantel tests: We computed raw Mantel tests using mantel.rtest in ade4 (Dray and Dufour 2007), specifying 1000 permutations to assess statistical significance. We computed phylogenetically permuted partial Mantel (pppMantel) tests (Lapointe and Garland 2001) using the script phyloMantel.R from Harmon & Glor (2010), again using 1000 permutations to test for significance.

Phylogenetic linear mixed models (PLMMs): For the character displacement datasets, we fitted PLMMs using asreml-R (Butler et al. 2009), including both sympatry and the patristic distance (calculated using the cophenetic.phylo function in ape [Paradis 2011]) between each pairwise species comparison as fixed effects and including the identity of each lineage and the phylogeny as random effects, following Tobias *et al.* (2014). To assess statistical significance of the fixed effects, we used Wald-type F-tests with Kenward-Rogers adjustments to the denominator degrees of freedom using the wald.asreml function with the option denDF = “numeric” in asreml-R, again following Tobias *et al.* (2014)

For the scenario using traits to predict species interactions, we fit PLMMs using MCMCglmm in R (Hadfield 2010), because the residual maximum likelihood approach used in asreml-R may bias estimates for logistic regressions (Bolker et al. 2009). We used standard inverse-Gamma priors for the fixed and random effects, using the code:

prior**<-list(**G**=list(**G1**=list(**V**=1,**nu**=0.002),**G2**=list(**V**=1,**nu**=0.002),**G3**=list(**V**=1,**nu**=0.002)),**R**=list(**V**=1,**nu**=0.02))**

We ran each fit for between 2 million and 20 million chains based on preliminary assessment of convergence for different tree sizes, varying the burn-in and thinning periods to result in approximately 2000 runs. We visually inspected convergence for a large sample of model fits to make sure that our MCMC parameterization was working well. MCMC fits were very computationally expensive, so we fit the models to a subset of simulated datasets (*n* = 100 per tree size, biogeography, and simulation parameter combination).

For all PLMMs, we randomized the order in which the identity of the lineages was passed to the random effects.

Simulation approach: BM models were first fit to simulated trait data using mvBM in the mvMORPH package (Clavel et al. 2015). Then, 5,000 datasets were simulated using the maximum likelihood estimate of the BM rate parameter and state at the root using fastBM in phytools (Revell 2012). A GLM (as outlined in “non-phylogenetic regressions”, the predictor variable for character displacement analyses was biogeographical overlap and the response variable was pairwise trait similarity, and for analyses of species interactions, the pairwise trait similarity was the predictor variable and a binomial variable of species interaction was the response variable) was fit to each of these simulated datasets, and the resulting analysis was considered statistically significant if both (a) the non-phylogenetic test on the raw data was statistically significant and (b) the test statistic from the raw analysis was outside of the 2.5-97.5% quantile interval of test statistics estimated on the simulated datasets.

Sister taxa analyses: for trees of size 150 and larger, we culled sister taxa from the full trees. We then ran non-phylogenetic regressions, PLMMs, the simulation approach and sister taxa GLMs on the culled dataset.

Phenotypic models: We fit all phenotypic models by maximum likelihood. We fit BM and OU models using geiger (Pennell et al. 2014) and the matching competition and two diversity dependent models (with either linear or exponential dependence of sigma on the number of species) using RPANDA (Morlon et al. 2016) in R. For the latter models, we included the biogeography used to simulate the datasets in the model fits, as described in Drury *et al.* (2016). We then compared the relative support of these models using Akaike weights (Burnham and Anderson 2002), since the models are not nested. Since model-fitting is computationally expensive, we fit the process-based models to a subset of datasets simulated on 100 tip trees (*n* = 100 biogeography and simulation parameter combination).

Additionally, to analyze the effect of *m* and *ψ* on rates of evolution, we fit four trait models to the sister-taxa datasets for both divergent and convergent character displacement scenarios using the R package EvoRAG (Weir and Lawson 2015): BM, OU, plus versions of BM and OU that allow the rate of trait evolution to change with the number of sympatric lineages (i.e., “BM_null”, “OU_null”, “BM_linear”, and “OU_linear_beta”).

---

**Supplementary Diagram 1.** Analytical methods used for each simulation scenario (see Box 1 for more details).

| analytical method | divergent character displacement |  | convergent character displacement |  | predicting spp. interactions |
| --- | --- | --- | --- | --- | --- |
|  | whole tree | sister taxa | whole tree | sister taxa | sympatric spp. |
| (1) non-phylogenetic regression | ✓ | ✓ | ✓ | ✓ | ✓ |
| (2) Mantel test | ✓ |  | ✓ |  |  |
| (3) phylogenetically permuted partial Mantel tests | ✓ |  | ✓ |  |  |
| (4) phylogenetic linear mixed models | ✓ | ✓ | ✓ | ✓ | ✓ |
| (5) phylogenetic simulations | ✓ | ✓ | ✓ | ✓ | ✓ |
| (6) process-based models using maximum likelihood | ✓ |  |  |  |  |
| (7) sister taxa GLMs |  | ✓ |  | ✓ |  |
| (8) sister taxa process-based models in EvoRAG |  | ✓ |  | ✓ |  |

**Supplementary Figure 1.**
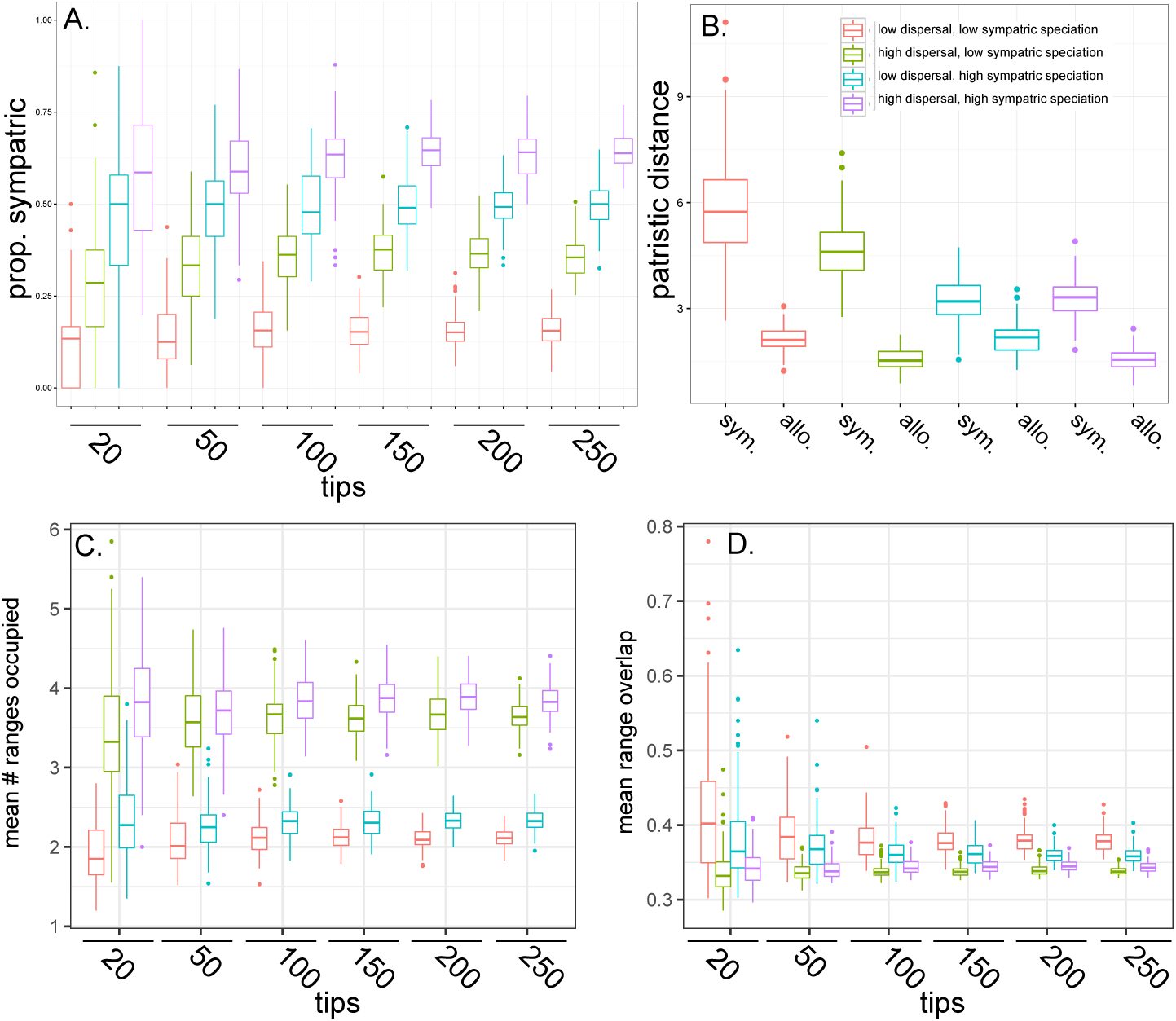
The resulting biogeographic landscape under each of the four biogeographic scenarios. A. The proportion of sister taxa that are living in sympatry varies as a function of both the dispersal rate and the level of sympatric speciation. B. The patristic distances (the branch lengths separating sister taxa) are larger for sympatric taxa, although the magnitude of this difference depends on the biogeographic scenario. C. The mean number of ranges occupied at the tips is higher under high dispersal rates. D. The mean range overlap for sympatric species pairs, calculated as the number of areas two species both occur in, divided by the total number of areas occupied by the lineage with the smaller range.

**Supplementary Figure 2.**
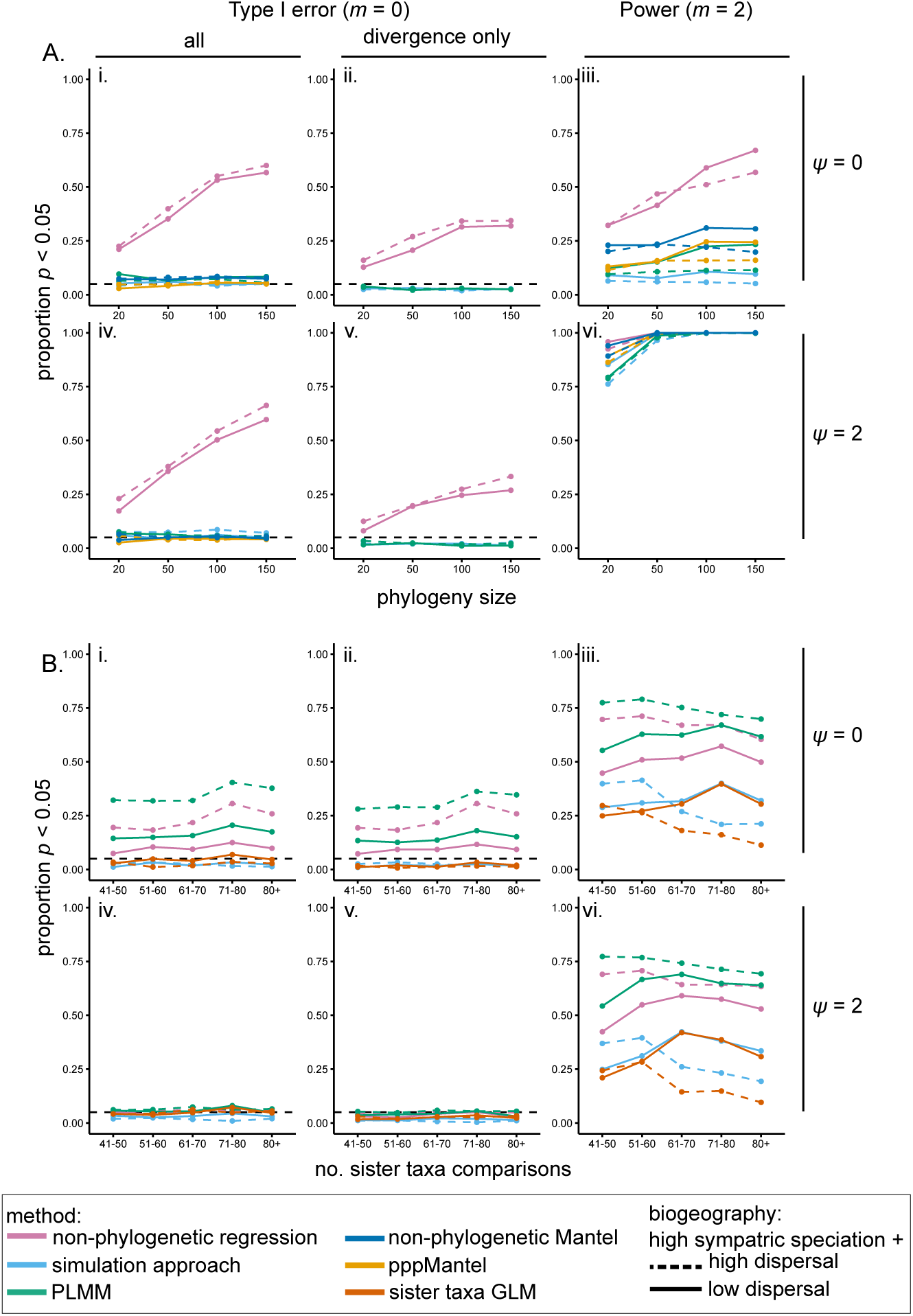
Proportion of statistically significant analyses in datasets simulated under divergent character displacement in biogeographic scenarios with high sympatric speciation rates. A. Results from approaches using data from all pairwise comparisons in a clade, plotted as a function of the phylogeny size and dispersal rate when i-ii. *m* = 0 and *ψ* = 0 (i. all analyses and ii. only analyses returning divergence in sympatry), iii. *m* = 2 and *ψ* = 0, iv-v. *m* = 0 and *ψ* = 2 (iv. all analyses and v. only analyses returning divergence in sympatry), and vi. *m* = 2 and *ψ* = 2. B. Results from analyses of sister-taxa culled from complete phylogenies binned by the number of resulting species pairs, plotted as a function of the number of sister taxa comparisons and dispersal rate when i-ii. *m* = 0 and *ψ* = 0 (i. all analyses and ii. only analyses returning divergence in sympatry), iii. *m* = 2 and *ψ* = 0, iv-v. *m* = 0 and *ψ* = 2 (iv. all analyses and v. only analyses returning divergence in sympatry), and vi. *m* = 2 and *ψ* = 2. For scenarios where *m* = 2, only the proportion of significant results showing divergence are plotted. Dashed horizontal lines represent a Type I error rate of 5%.

**Supplementary Figure 3.**
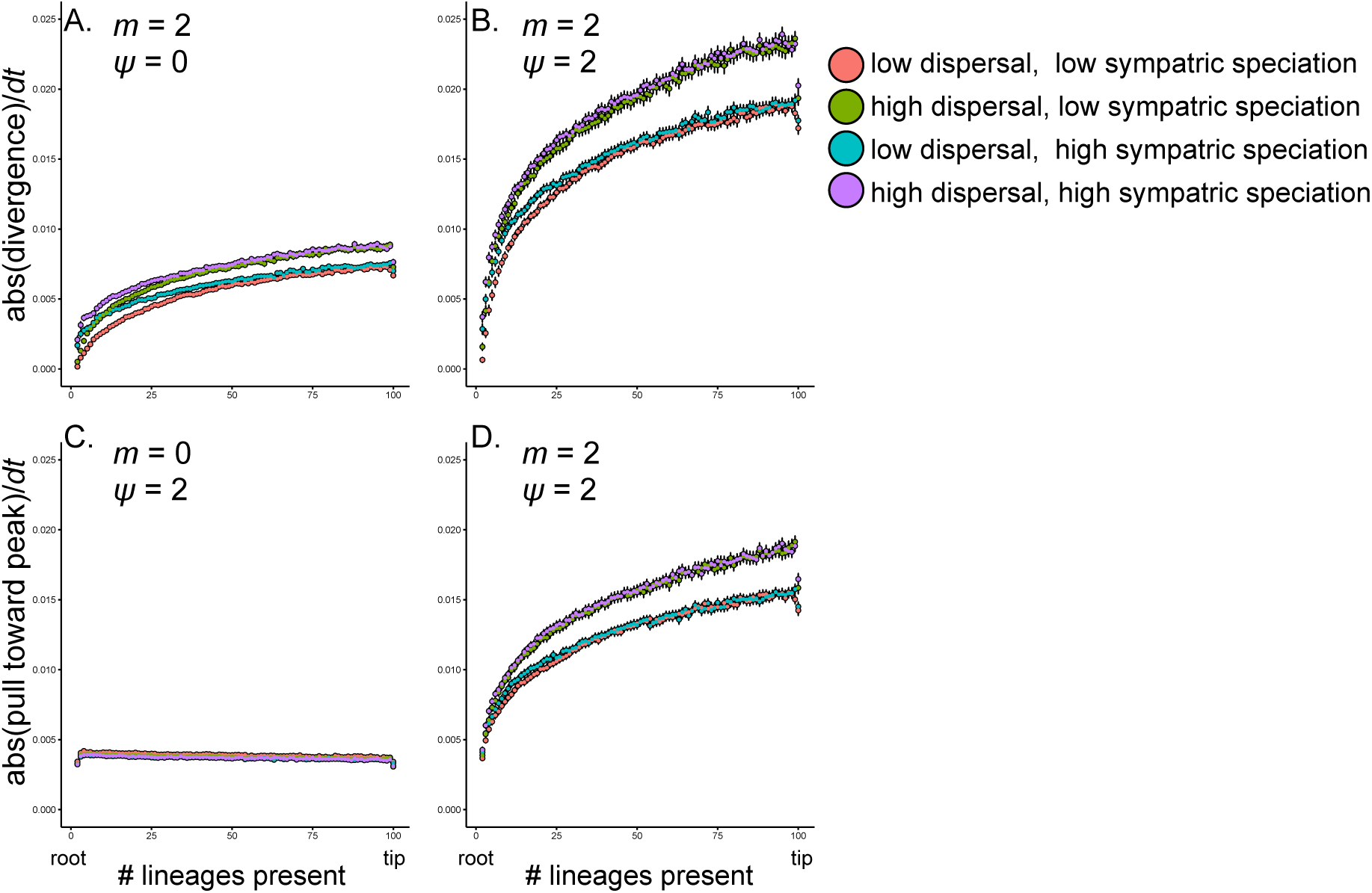
The effect of repulsion on the magnitude of changes in traits from the root to the tip of phylogenies under different simulation scenarios. To calculate the absolute value of divergence steps (A & B), we recorded the absolute value of the magnitude of the step size resulting from the deterministic repulsion step of Eq. 1 (i.e., 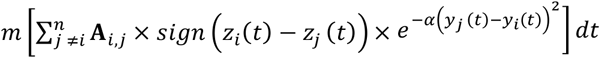 at each time step, took the mean during each internode interval for all extant lineages within trees, and then averaged each internode interval across all simulations. Each point thus represents the mean of means and the standard error across 100 phylogenies. The absolute value of the steps pulling trait values toward a stable peak (C & D) similarly represents the absolute value of the magnitude of the step size resulting from the deterministic component of the OU attraction to the peak in Eq. 1 (i.e., *ψ*(*θ* − *z_i_(t))dt*).

**Supplementary Figure 4.**
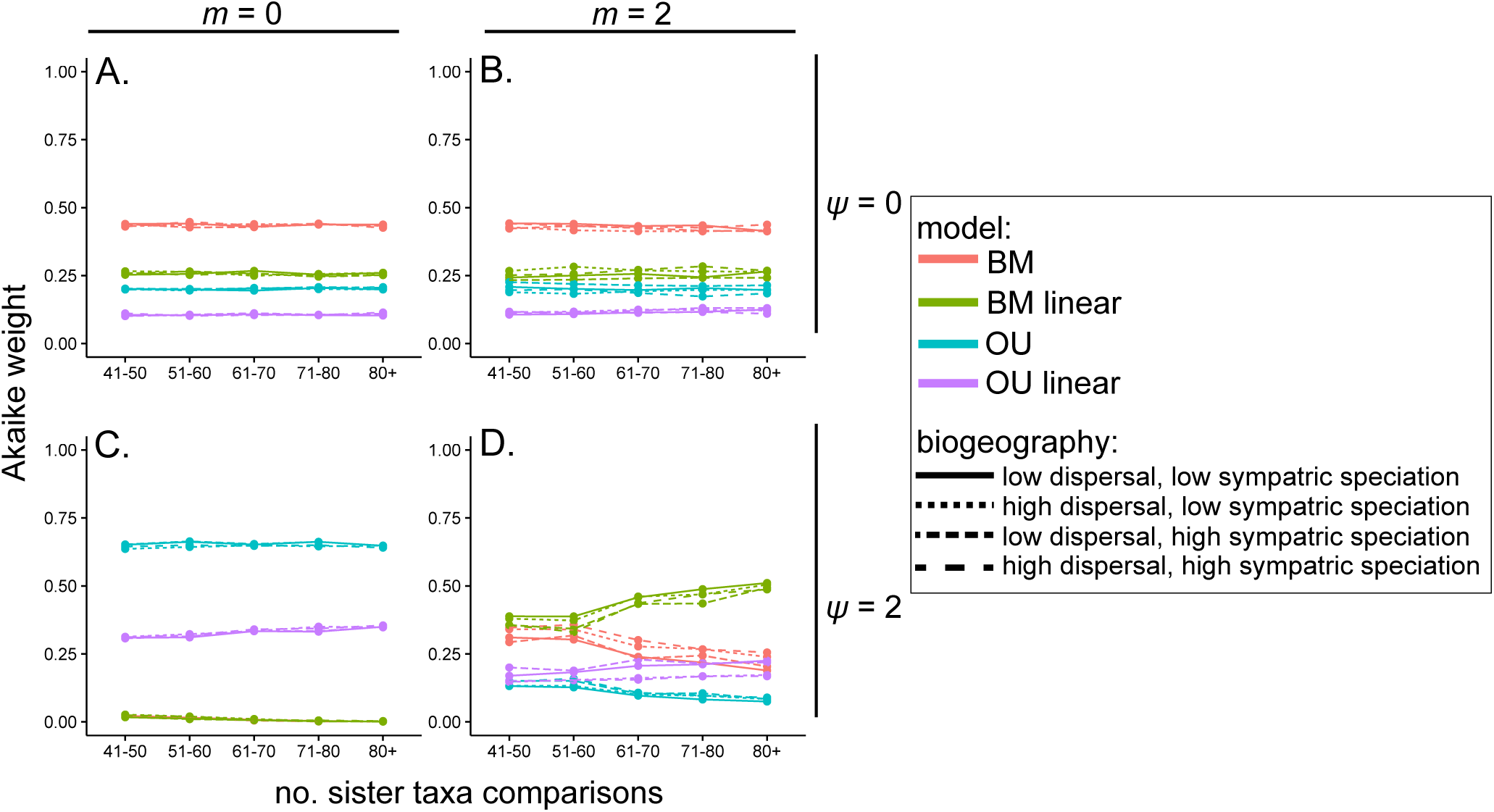
Models fit to sister-taxa datasets generated under divergent character displacement scenarios in EvoRAG corroborate the influence of the number of sympatric lineages on the rate of evolution. A. When *ψ =* 0, BM is the best fitting model, whether A. *m* = 0 or B. *m* = 2. When *ψ =* 2, OU is the best fitting model when C. *m* = 0, whereas a BM model that allows the rate of trait evolution to vary linearly with the number of sympatric lineages is a best fitting model when D. *m* = 2. Plots depict the Akaike weight of each model as a function of both the biogeographic scenario and the number of sister taxa comparisons.

**Supplementary Figure 5.**
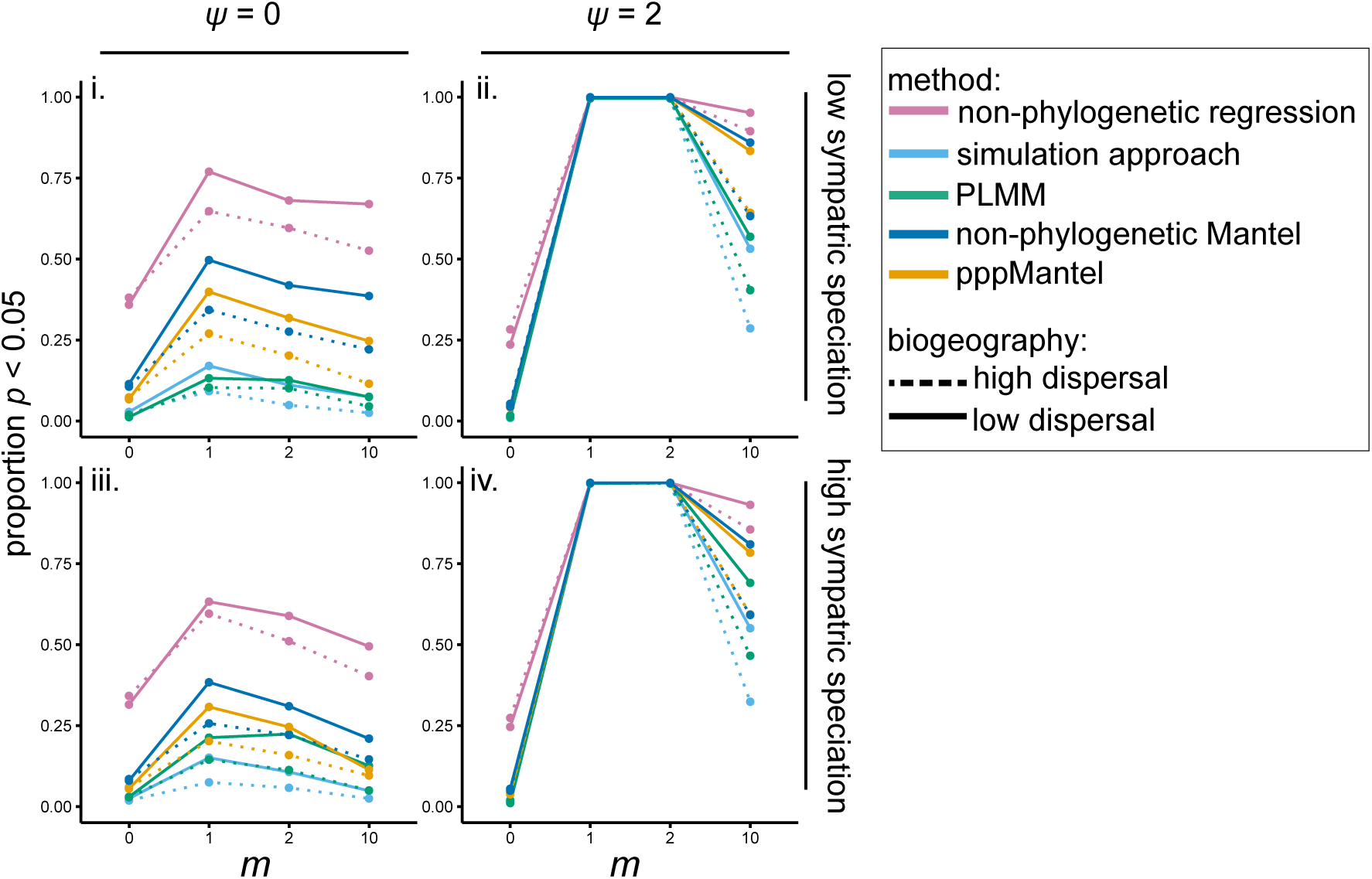
The proportion of statistically significant analyses for datasets with divergent character displacement varies as a function of both *m* and *ψ.*

**Supplementary Figure 6.**
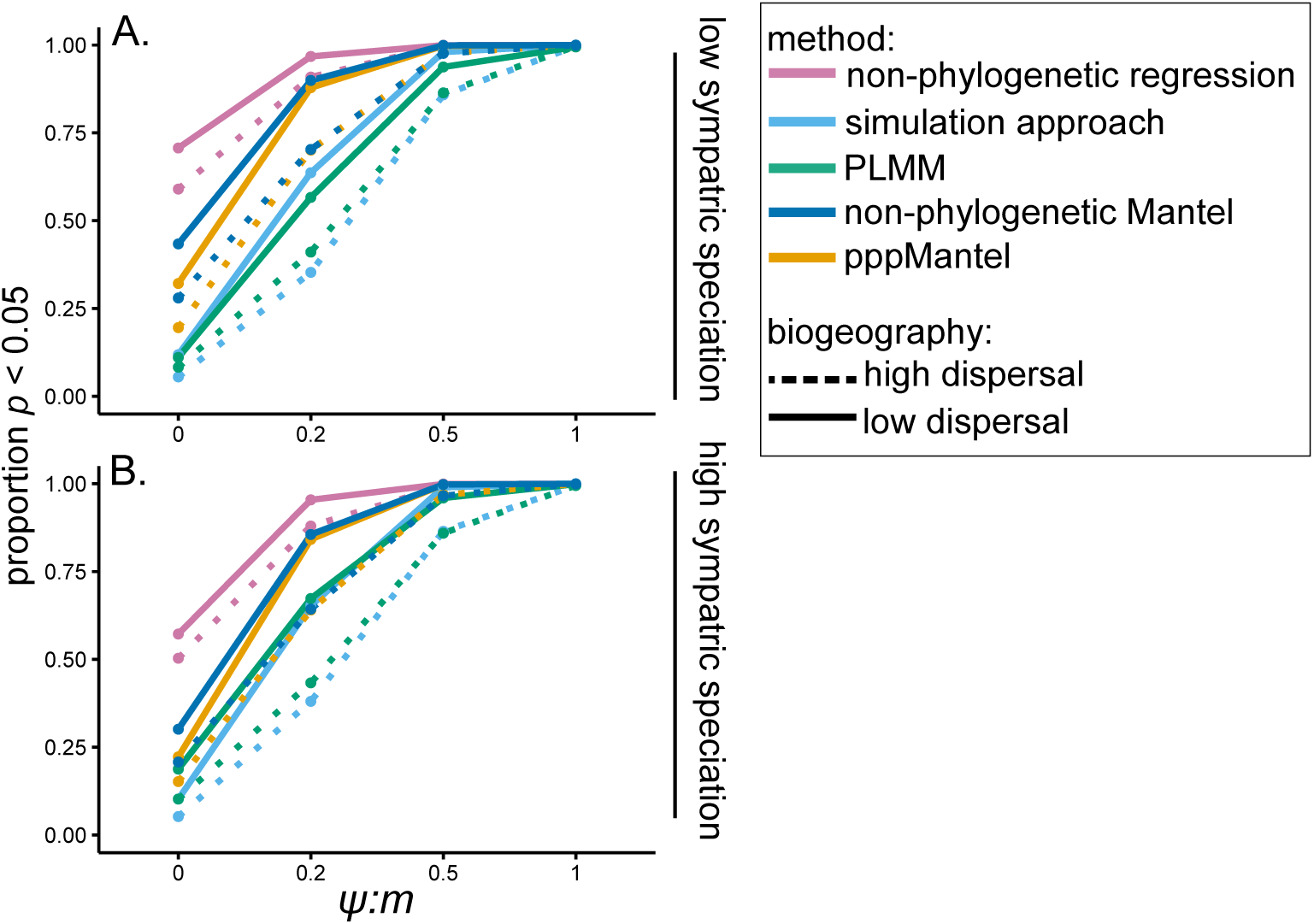
As *ψ:m* increases, the ability of all methods to detect character displacement, when present, increases under both low sympatric speciation (A) and high sympatric speciation (B) scenarios.

**Supplementary Figure 7.**
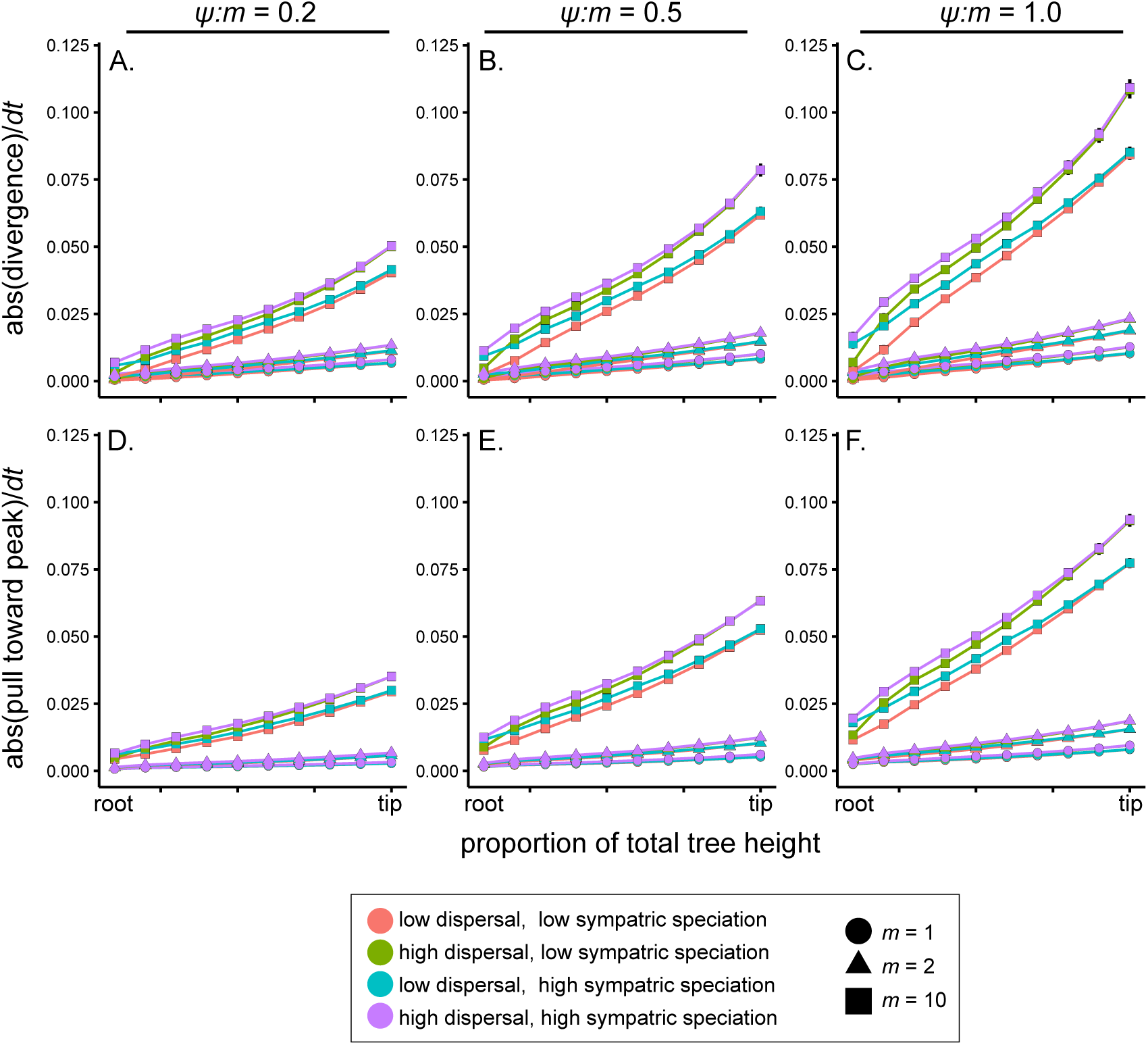
The effect of repulsion on the magnitude of changes in traits from the root to the tip of phylogenies as a function of the ratio *ψ:m.* As *ψ* increases relative to m, both the magnitude of divergence steps (panels A-C) and the magnitude of the steps pulling trait values toward the peak (panels D-F) increase. This effect becomes increasingly apparent at higher *m* values and in biogeographic scenarios generated under high dispersal rates. For a description of the data, see the legend of Fig. S3. In this plot, data are averaged within 10 time bins representing each 10% of the total height of the phylogeny.

**Supplementary Figure 8.**
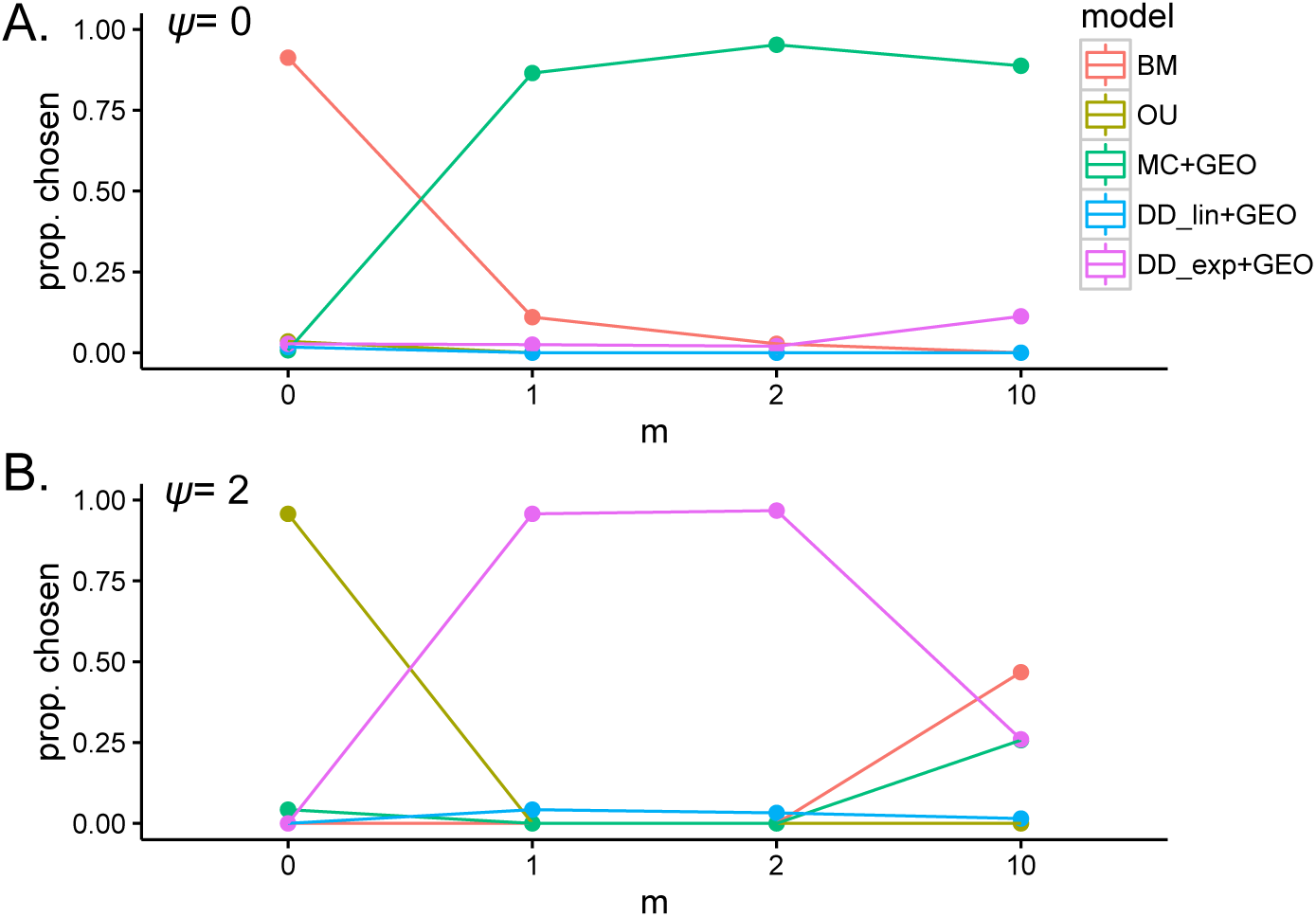
Proportion of simulated datasets for which a given phenotypic model was chosen using model selection as a function of *m*. We considered a model to be chosen if it had the lowest AICc score among all of the models, and if the AICc score was > 2 units away from BM model. A. Results for datasets simulated without the OU process. B. Results from datasets simulated with *ψ* = 2.

**Supplementary Figure 9.**
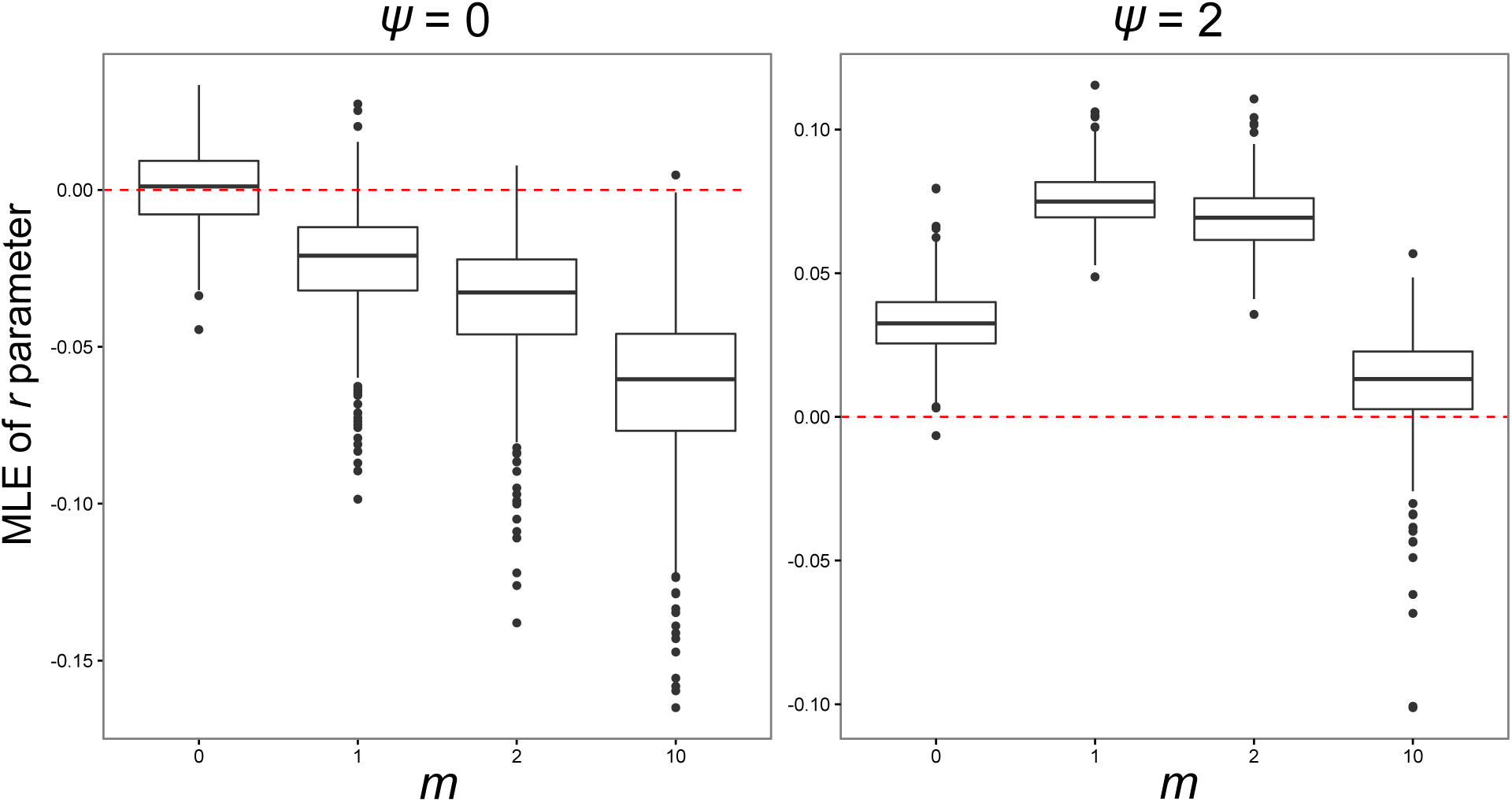
The rate parameter *r* of the DDexp+GEO model is increasingly negative with increasing *m* values when *ψ* = 0, yet is positive when *ψ* > 0.

**Supplementary Figure 10.**
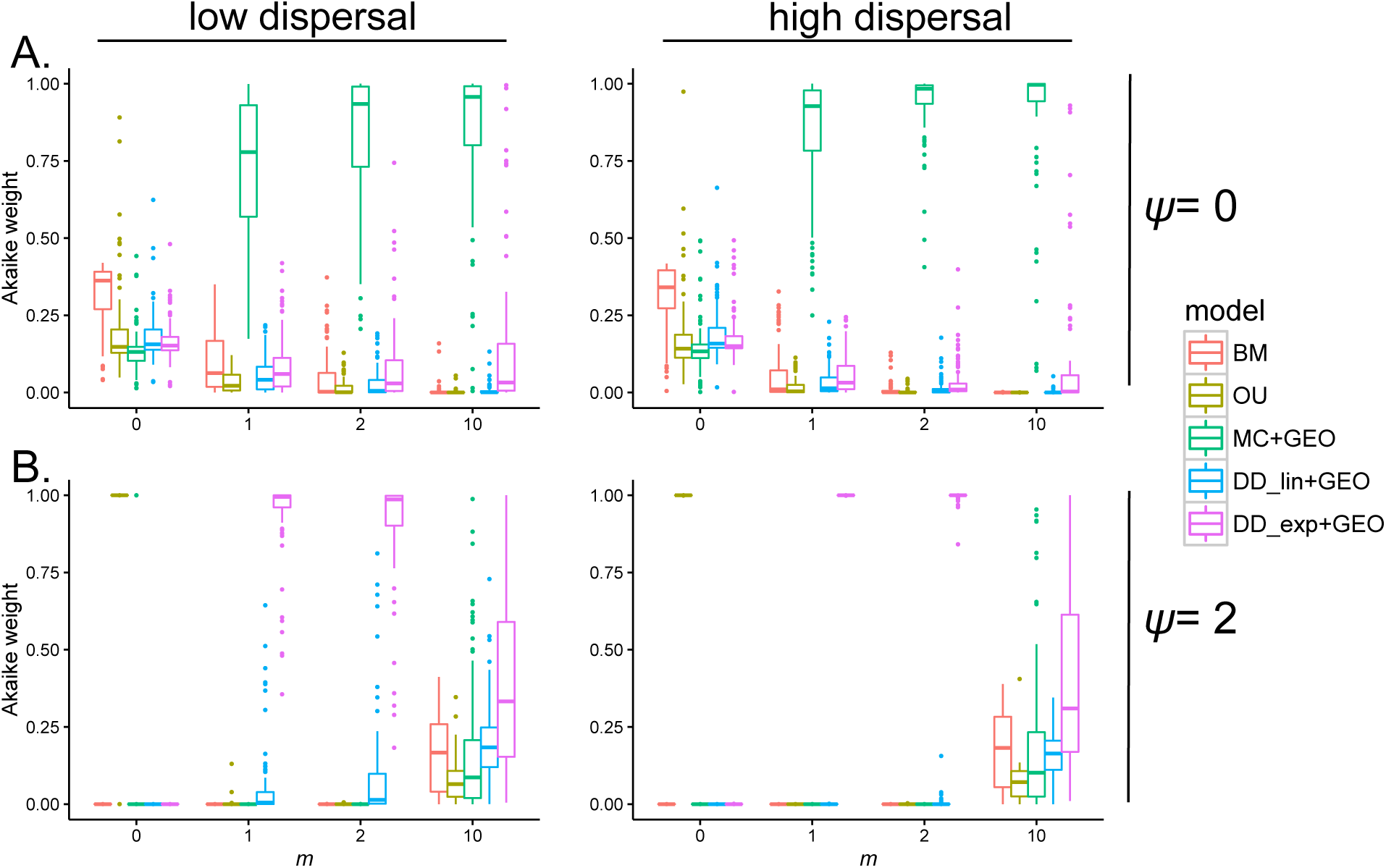
Akaike weights for each trait model fit to simulated datasets in biogeographic scenarios with high sympatric speciation rates as a function of *m.* A. When OU is absent, BM is the best-fit model when *m* = 0, and the matching competition model with biogeography is the best model when competitive divergence is present. B. When OU is present, OU is the best-fit model when *m* = 0, and the diversity-dependent exponential model with biogeography is the best model when competitive divergence is present and *ψ:m* is relatively high.

**Supplementary Figure 11.**
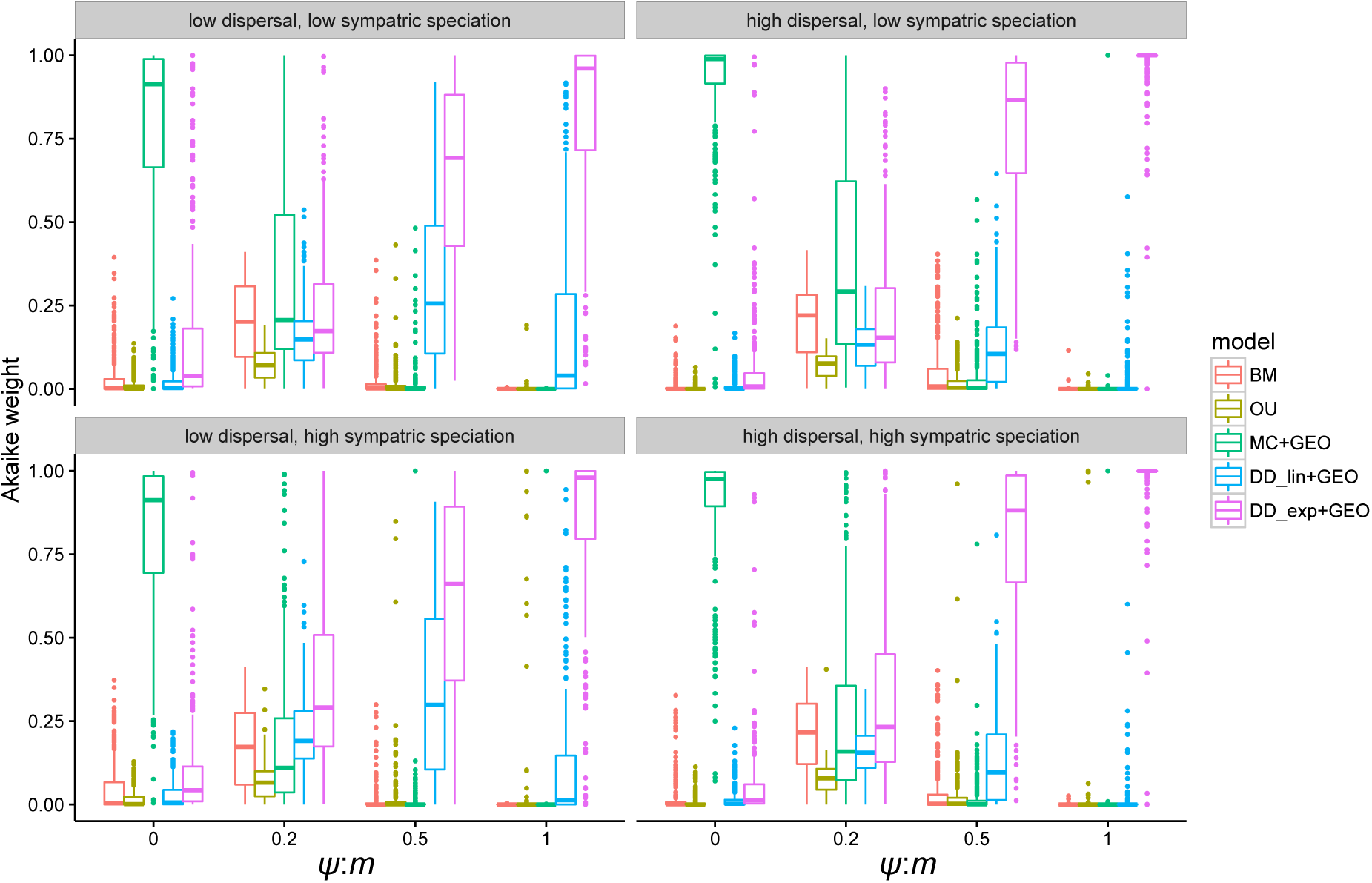
As *ψ:m* increases, the DDexp + GEO model is increasingly better fit to datasets simulated with character displacement.

**Supplementary Figure 12.**
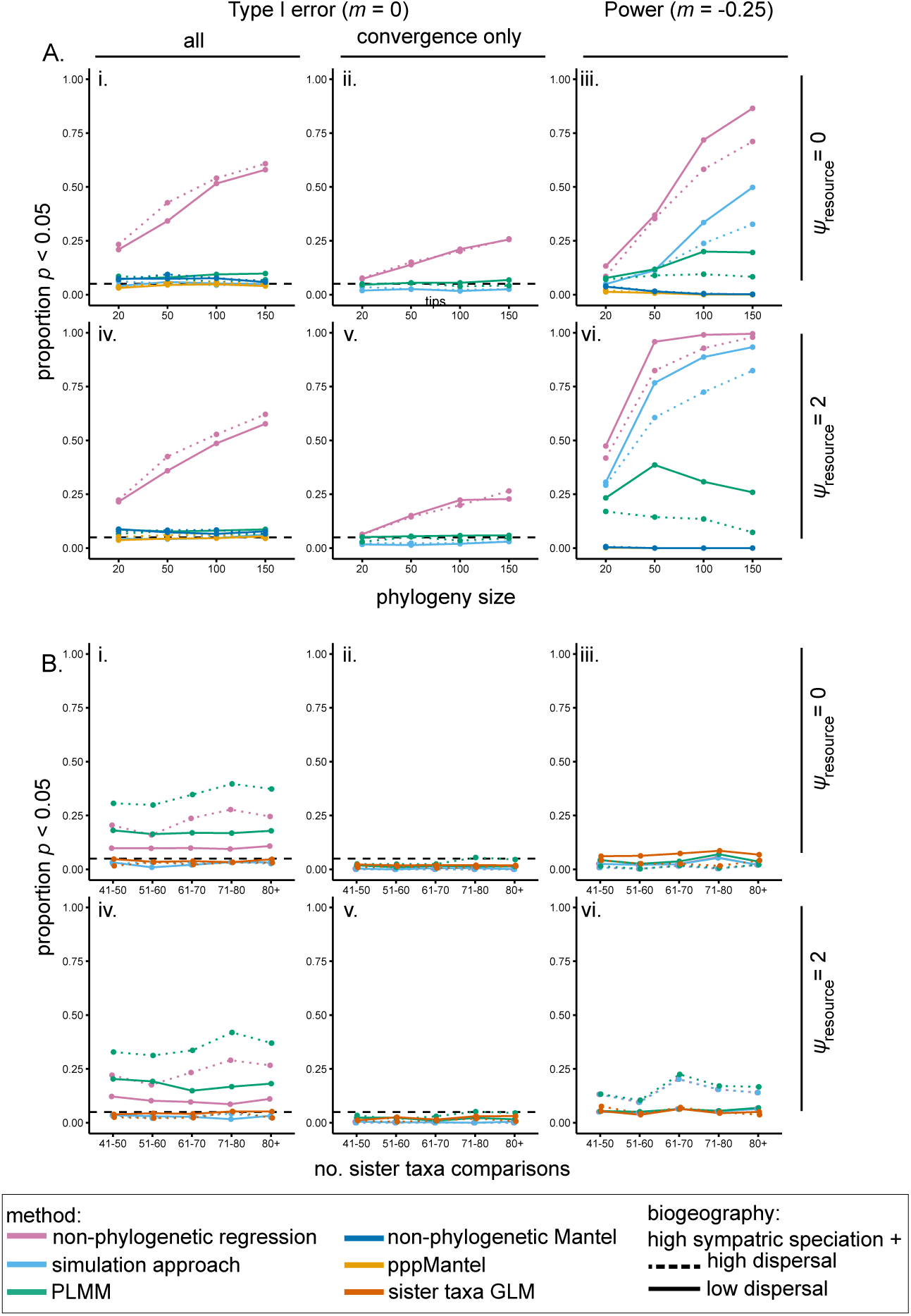
Proportion of statistically significant analyses in datasets simulated under convergent character displacement in biogeographic scenarios with high sympatric speciation rates. A. Results from approaches using data from all pairwise comparisons in a clade, plotted as a function of the phylogeny size and dispersal rate when i-ii. *m* = 0 and *ψ_resource_*= 0 (i. all analyses and ii. only analyses returning convergence in sympatry), iii. *m* = −0.25 and *ψ_resource_* = 0, iv-v. *m* = 0 and *ψ_resource_* = 2 (iv. all analyses and v. only analyses returning convergence in sympatry), and vi. *m* = −0.25 and *ψ_resource_* = 2. B. Results from analyses of sister-taxa culled from complete phylogenies binned by the number of resulting species pairs, plotted as a function of the number of sister taxa comparisons and dispersal rate when i-ii. *m* = 0 and *ψ_resource_* = 0 (i. all analyses and ii. only analyses returning convergence in sympatry), iii. *m* = −0.25 and *ψ_resource_* = 0, iv-v. *m* = 0 and *ψ_resource_* = 2 (iv. all analyses and v. only analyses returning convergence in sympatry), and vi. *m* = −0.25 and *ψ_resource_* = 2. For scenarios where *m* = −0.25, only the proportion of significant results showing convergence are plotted. Dashed horizontal lines represent a Type I error rate of 5%.

**Supplementary Figure 13.**
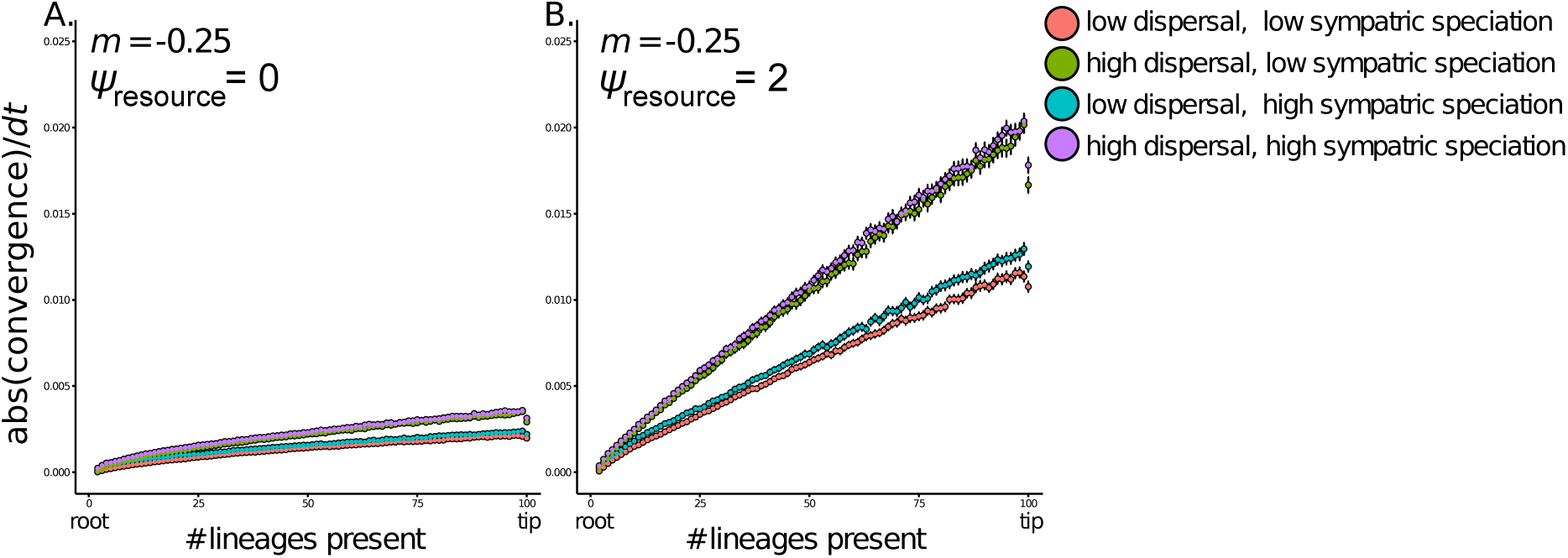
The effect of convergence on the magnitude of changes in traits in simulations, plotted as a function of stabilizing selection in the resource-use trait (*ψ*_resource_) and biogeography. With *ψ*_resource_ = 2, the magnitude of convergence steps is higher, and this is particularly true for high dispersal biogeographies. Data generated as described in the legend of Fig. S3.

**Supplementary Figure 14.**
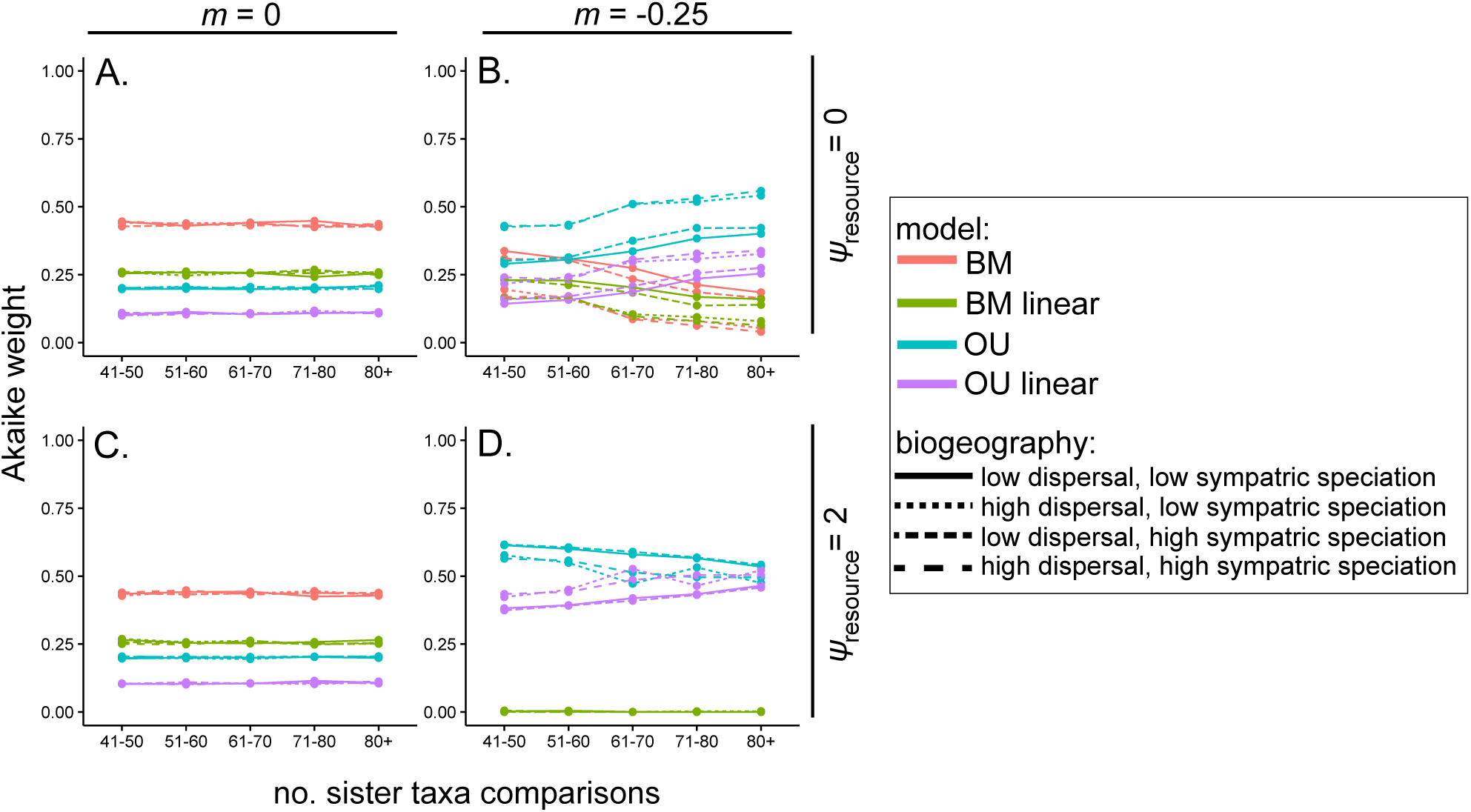
Models fit to sister-taxa datasets generated under convergent character displacement scenarios in EvoRAG. A. When *m* = 0, BM is the best fitting model, whether A. *ψ_resource_* = 0 or C. *ψ_resource_* = 2. When *m* = −0.25, OU is the best fitting model when C. *ψ_resource_* = 0 and D. *ψ_resource_* = 2, but an OU model with evolutionary rates that change linearly as a function of the number of sympatric lineages (OU linear) is also a strong model. Plots depict the Akaike weight of each model as a function of both the biogeographic scenario and the number of sister taxa comparisons.

**Supplementary Figure 15.**
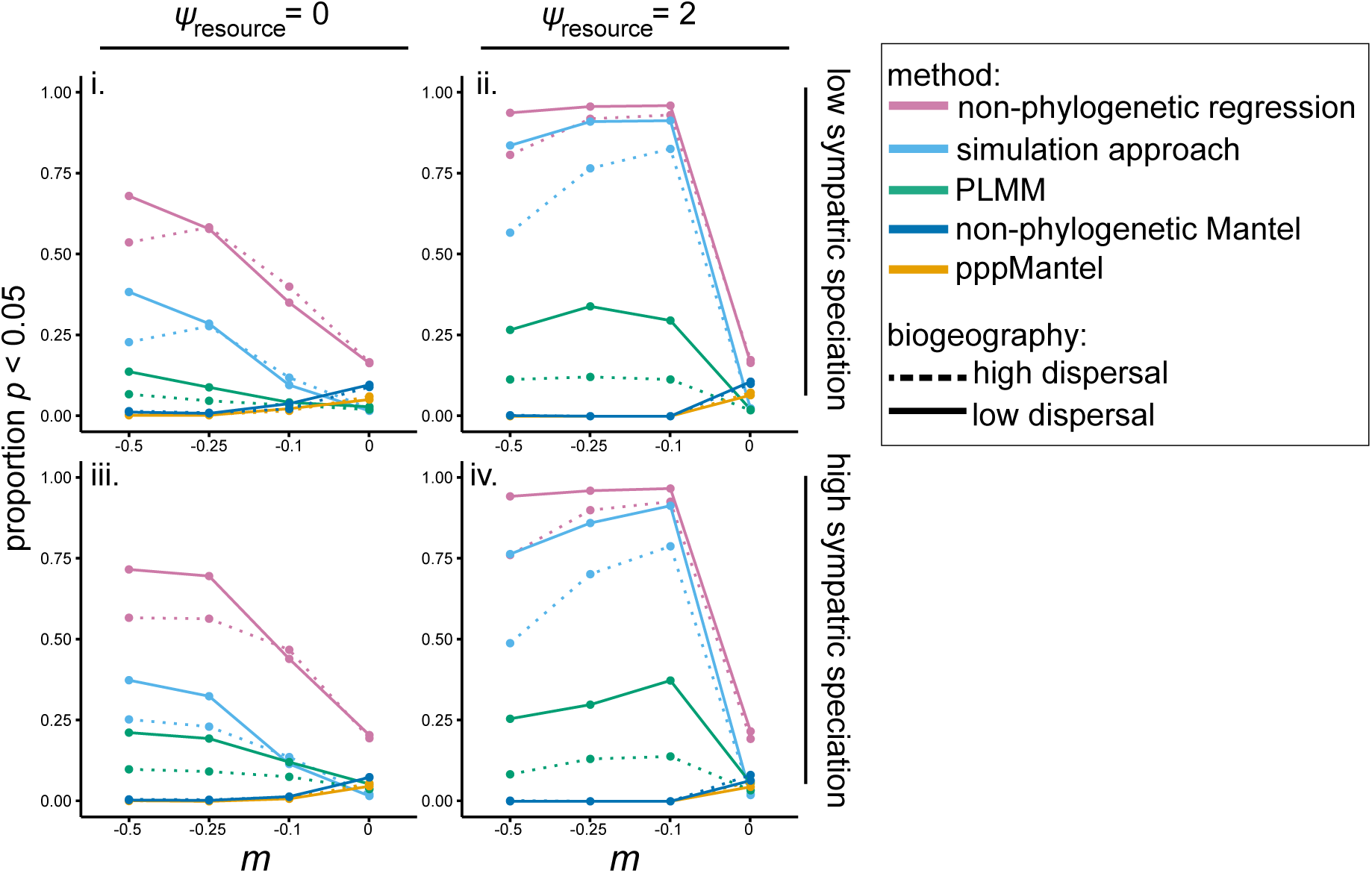
The proportion of statistically significant analyses for datasets with convergent character displacement varies as a function of both m, and is generally lower in the absence of a pull toward a peak (i,iii) compared to cases where data were simulated with a pull toward a peak (ii,iv).

**Supplementary Figure 16.**
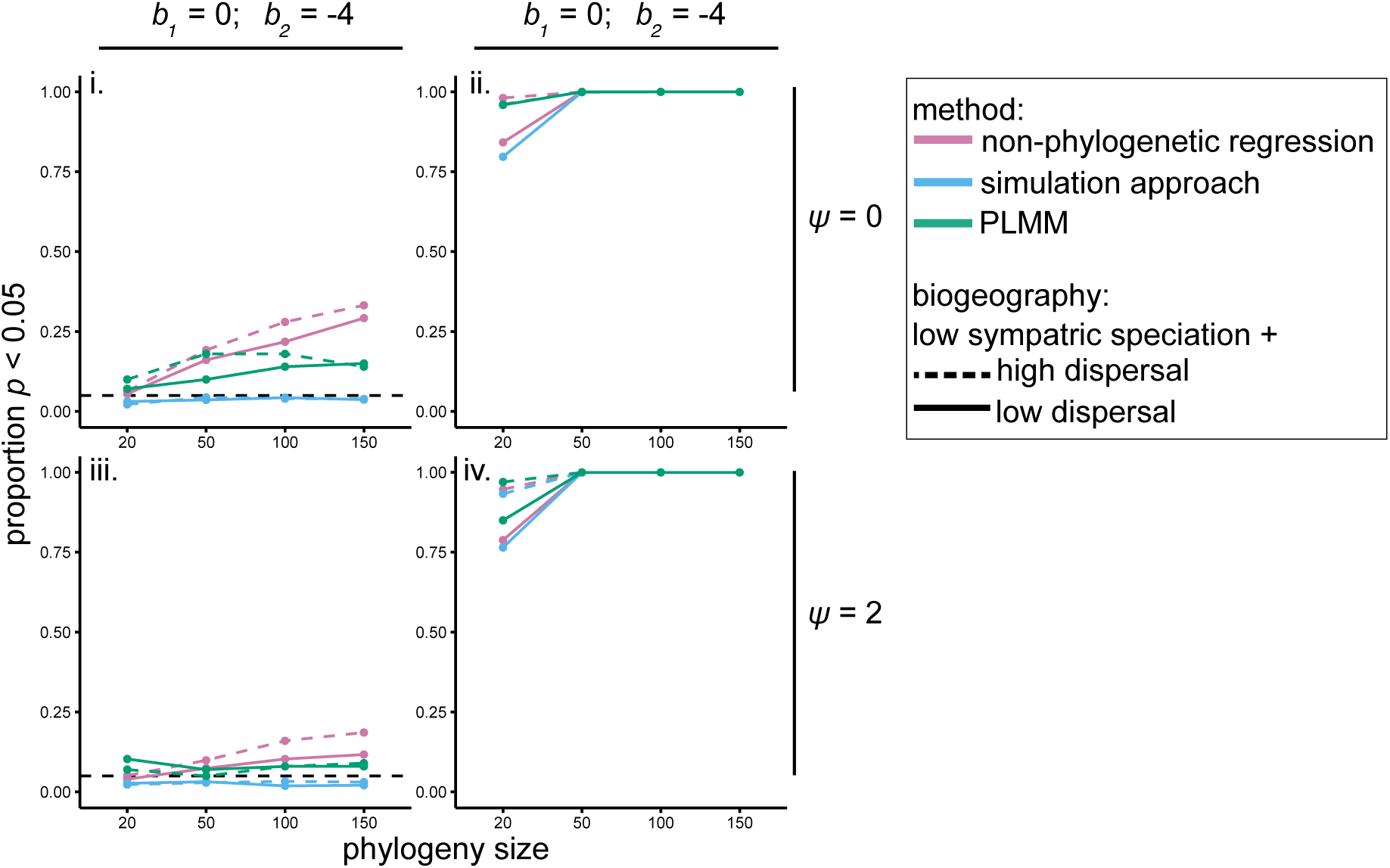
Proportion of statistically significant analyses in datasets with interactions simulated under a simple phenotype matching process in biogeographic scenarios with high sympatric speciation rates. Results from analyses where the focal trait was simulated under A. BM or B. OU, plotted as a function of the dispersal rate when i. *b_1_* (the simulation coefficient determining the relationship between the interaction and the measured trait) = 0, *b_2_* (the simulation coefficient for an unmeasured trait) = −4, and *ψ* = 2, ii. *b_1_* = −4, *b_2_* = 0, and *ψ* = 2, iii. *b_1_* = 0, *b_2_* = −4, and *ψ* = 0, and iv. *b_1_* = −4, *b_2_* = 0, and *ψ* = 0.

**Supplementary Figure 17.**
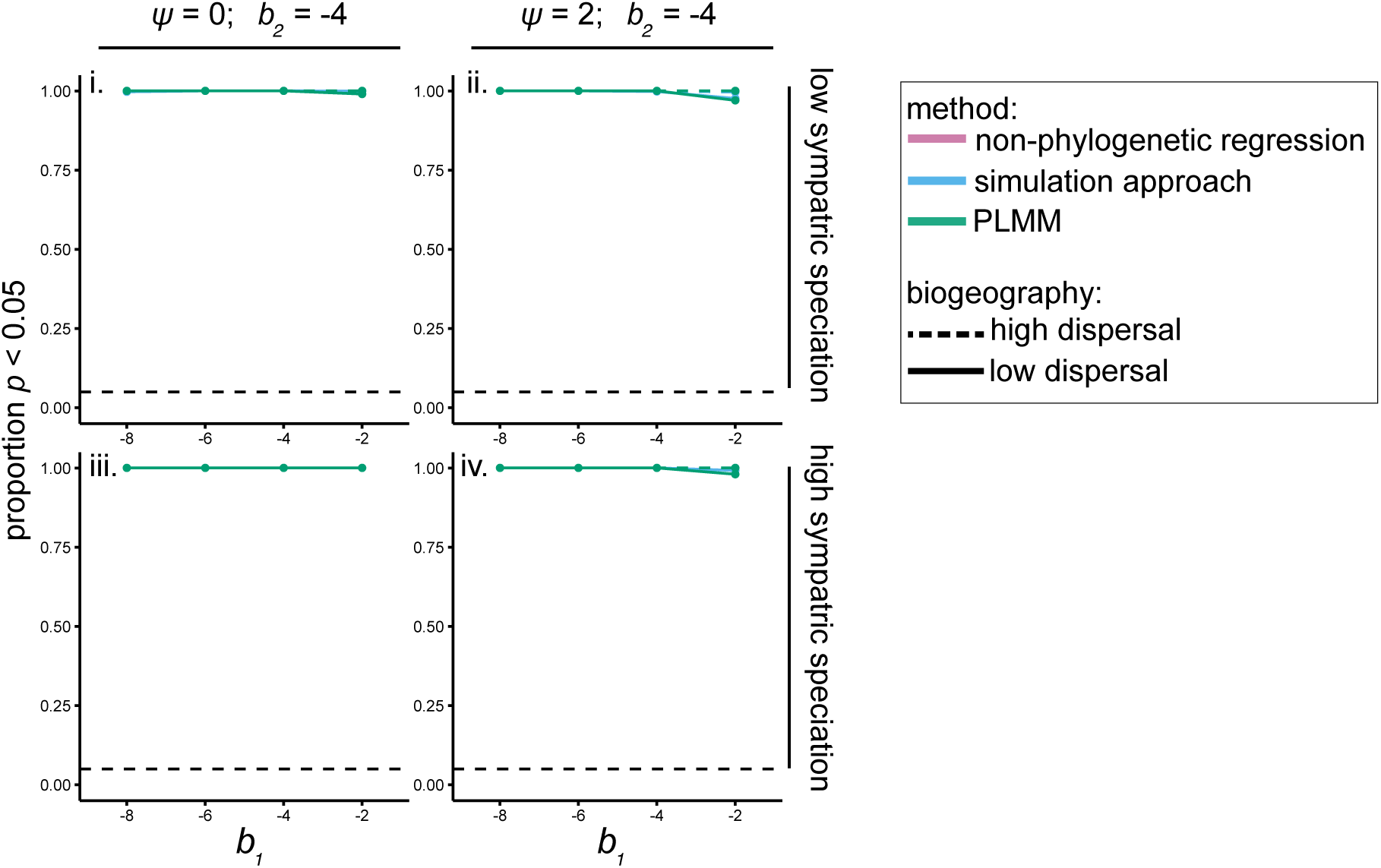
The power to detect a causal relationship between a species interaction and similarity in a trait value is not affected greatly by varying the coefficient determining the strength of this relationship, regardless of whether the OU process is absent (i, iii) or present (ii, iv) in the measured trait. In these simulations, tree size was held constant at 100 species.

